# Mining bacterial NGS data vastly expands the complete genomes of temperate phages

**DOI:** 10.1101/2021.07.15.452192

**Authors:** Xianglilan Zhang, Ruohan Wang, Xiangcheng Xie, Yunjia Hu, Jianping Wang, Qiang Sun, Xikang Feng, Shanwei Tong, Yujun Cui, Mengyao Wang, Shixiang Zhai, Qi Niu, Fangyi Wang, Andrew M. Kropinski, Xiaofang Jiang, Shaoliang Peng, Shuaicheng Li, Yigang Tong

## Abstract

Temperate phages (active prophages induced from bacteria) help control pathogenicity, modulate community structure, and maintain gut homeostasis^1^. Complete phage genome sequences are indispensable for understanding phage biology. Traditional plaque techniques are inapplicable to temperate phages due to the lysogenicity of these phages, which curb the identification and characterization of temperate phages. Existing in silico tools for prophage prediction usually fail to detect accurate and complete temperate phage genomes^2–5^. In this study, by a novel computational method mining both the integrated active prophages and their spontaneously induced forms (temperate phages), we obtained 192,326 complete temperate phage genomes from bacterial next-generation sequencing (NGS) data, hence expanded the existing number of complete temperate phage genomes by more than 100-fold. The reliability of our method was validated by wet-lab experiments. The experiments demonstrated that our method can accurately determine the complete genome sequences of the temperate phages, with exact flanking sites (*attP* and *attB* sites), outperforming other state-of-the-art prophage prediction methods. Our analysis indicates that temperate phages are likely to function in the evolution of microbes by 1) cross-infecting different bacterial host species; 2) transferring antibiotic resistance and virulence genes; and 3) interacting with hosts through restriction-modification and CRISPR/anti-CRISPR systems. This work provides a comprehensive complete temperate phage genome database and relevant information, which can serve as a valuable resource for phage research.

## Main Text

Temperate phage is a key component of the microbiome. Temperate phages have both lysogenic and lytic cycles. When they integrate (as prophages) into bacterial chromosomes, they enter the lysogenic cycle to participate in several bacterial cellular processes^6^. As the temperate phages have an inherent capacity to mediate the gene transfers between bacteria by specialized transduction, they may increase bacterial virulence and promote antibiotic resistance^1,7^. Therefore, temperate phage detection is necessary to ensure the safety of phage therapy. Also, isolating suitable virulent phages is challenging for some bacterial infections, such as *Clostridioides difficile* and *Mycobacterium tuberculosis*^8–10^. Hence, genetically modifying temperate phages into virulent phages is a promising phage therapy approach for such bacterial infections. In recent years, fecal microbiota transplantation (FMT) has increasingly become a prominent therapy to treat *C.difficile* infection (CDI) by normalizing microbial diversity and community structure in patients^11,12^. FMT may also help manage other disorders associated with gut microbiota alteration^11^. However, drug-resistant and highly virulent bacteria in the donor’s stool pose a potential threat to the patient’s life. Identifying and profiling the temperate phages that carry important antibiotic resistance genes and virulence genes, is crucial for FMT success.

Even though temperate phages play essential roles in medical treatments, only a few genomes are accessible untill now, seriously hindering the related research. Using integrase as a marker, we identified only 1,504 complete temperate phage genomes on GenBank (December 2020, Extended Data Table 4). Detecting temperate phages is challenging. Traditionally, phage detection relies on culture-based methods to isolate the temperate phage after inducing it from a lysogenic strain, amplify the phage to high titers, and characterize it often by transducing the phage into different strains^13^. However, as temperate phages are readily integrated into the host genome and stay silent, forming phage plaques is difficult after transduction, which causes a big challenge for the traditional culture method. Nowadays, an increasing amount of bacterial genomic sequences have become available in databases due to the advances in next-generation sequencing (NGS) technologies. Bioinformatics tools have been developed to predict potential prophage sequences within bacterial genomes, including Phage_Finder^14^, Prophage finder^15^, Prophinder^16^, PHAST^2^, PHASTER^17^, PhiSpy^3^, VirSorter^18^, Prophage Hunter^4^, and VIBRANT^5^. Among these tools, PHASTER/PHAST, VirSorter, and Prophage Hunter consider the completeness of a predicted prophage sequence. Typically, these tools rely on phage protein clusters to locate possible prophage regions on assembled bacterial genome sequences, yet fail to acquire exact active prophage (temperate phage) sequence boundaries^2–5^. That is, all the existing prophage prediction tools are unable to obtain accurately complete temperate phage genome sequences.

Here we present a novel computational method to acquire complete temperate phage genome sequences using bacterial NGS data. Unlike other in silico tools that adopt machine learning or statistical classifier to predict possible prophage regions from the bacterial genomes (Table 1), our method utilizes raw NGS data (reads) to identify sequences that match the circularized temperate phage genome. The raw reads capture the biological replication process signals of temperate phages in their lytic cycles. Essentially, our method incorporates three biological principles from phage life cycles to detect temperate phages: 1) In the replication process of the lytic cycle, the phage genome circularizes^19,20^. 2) During the cycle of lysogenization, a temperate phage integrates into the host chromosome and shares a core sequence of attachment sites (*attP*/*attB*) with its host genome sequence (Figure 1). 3) Active prophages usually undergo spontaneous induction at a relatively low frequency, resulting in a small amount of temperate phages in the bacterial culture, which will generate circularized phage genome’s reads in the bacterial NGS data. As we label a genome sequence as temperate phage only and only if 1) this genome sequence takes a part of its host’s assembled contig and 2) this genome is also circular, plasmid genome sequences are excluded.

**Figure 1.**
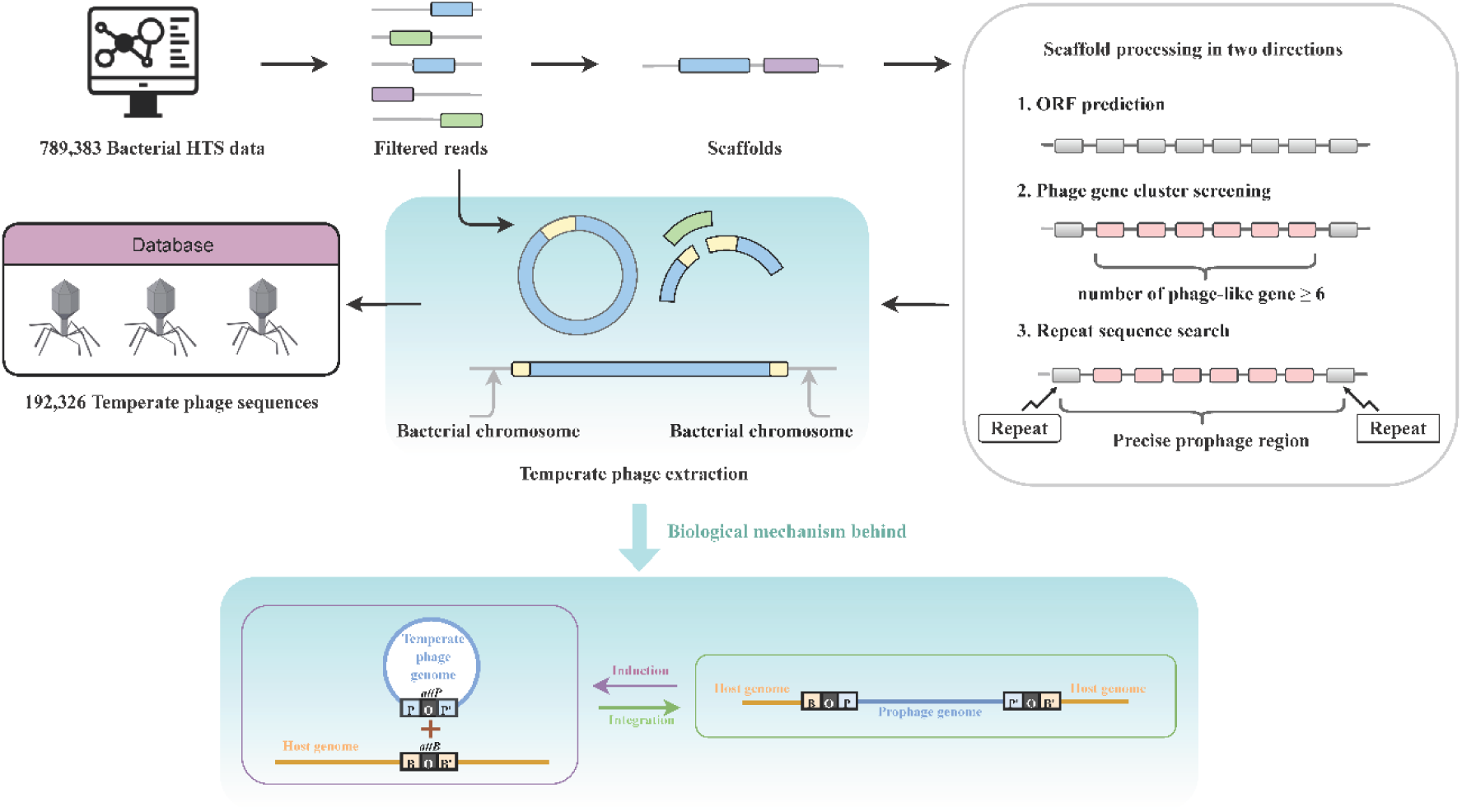
Workflow of our temperate phage detection method and illustration of temperate phage induction and integration processes. The main step of temperate phage detection is based on the temperate phage induction and integration processes, which is illustrated at the bottom of the figure. *attP* is short for attachment site of phage, *attB* represents the attachment site of host strain, *attL* stands for the left attachment site after integration, *attR* is short for the right attachment site after integration, O stands for core region of phage and bacterium, B represents host, while P represents phage. Temperate phage is also called prophage after integrating into a lysogenic host strain.

**Table 1.**
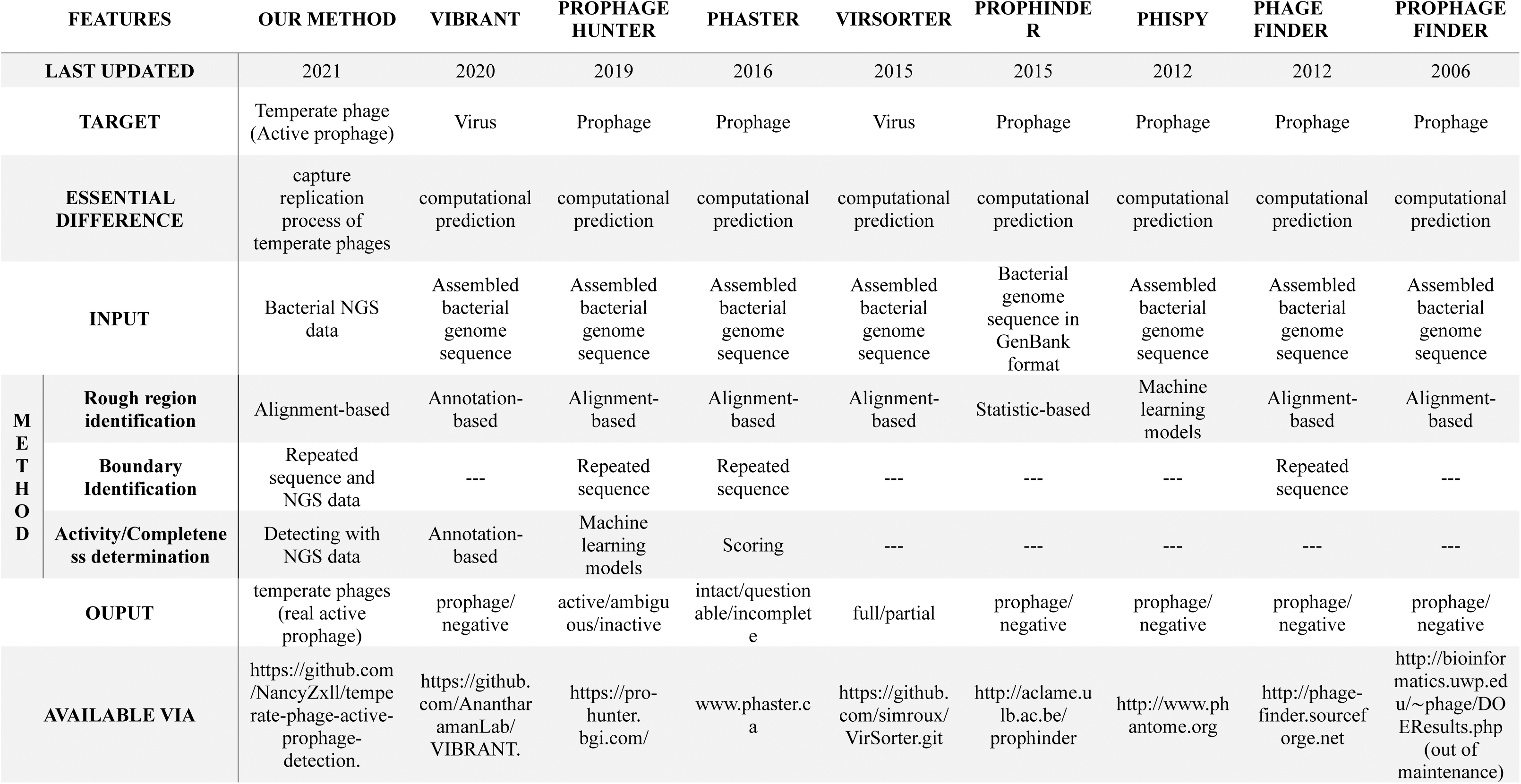
General characteristics of our method and eight commonly used prophage prediction tools.

Accordingly, the computational temperate phage detection method consists of three steps: NGS data processing, prophage region detection, and temperate phage extraction (Figure 1 and Methods). In principle, our method can detect all the temperate phages when they are spontaneously induced to a particular concentration from their host strains.

To validate our method, we sequenced the bacterial strains preserved in our laboratory using NGS technology (Extended Data Table 1). In total, our computational method detected 17 temperate phages from 15 bacterial strains. Subsequently, the wet-lab experiments were then conducted to induce the temperate phages from these bacterial strains. These wet-lab induced temperate phages were then sequenced using NGS technology, and their genome sequences were treated as ground truth. In contrast to Prophage finder^15^, Prophinder^16^, PHASTER^17^, PhiSpy^3^, VirSorter^18^, Prophage Hunter^4^, and VIBRANT^5^, our method identified exact boundaries and acquired accurate temperate phage genome sequences (Extended Data Figure 1, Extended Data Table 2 and Supplementary Materials, Verification of our temperate phage detection method).

After applying the filtration criteria on NGS data (Extended Data Table 5), we downloaded 789,383 raw NGS host data sets from GenBank (March 2020) and analyzed them using the proposed temperate phage detection method. We discovered 192,326 complete temperate phage sequences within 2,717 host species of 710 host genera (Extended Data Table 6, Supplementary Materials, Expansion of temperate phages). Among them, *Salmonella enterica* has the most temperate phages, followed by *Escherichia coli*, *Listeria monocytogenes*, *Klebsiella pneumoniae*, *Staphylococcus aureus*, etc. (Extended Data Figure 3C). The temperate phages in *L. monocytogenes* have the widest genome sizes, while these in human gut metagenome have the broadest range of GC content (Figure 2). By grouping the identical/reverse complementary sequences, we harvest 66,823 nonredundant complete temperate phage genomes in total.

**Figure 2.**
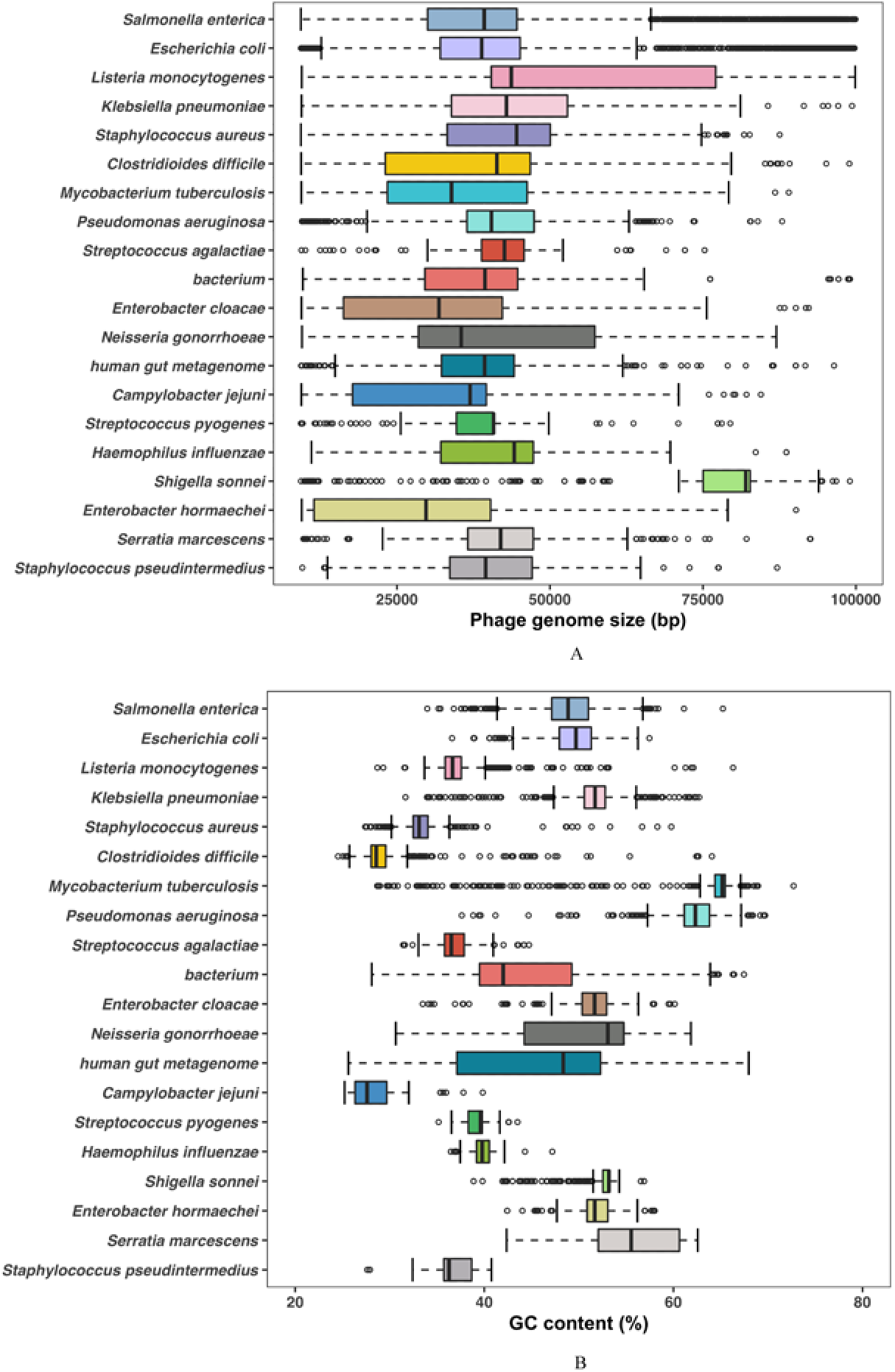
Genome size and GC content distribution of temperate phages in the top 20 host species. The distribution of genome size (A) and GC content (B) distribution in the top 20 most host species. The ’bacterium’ relates to the same name listed in the species of GenBank taxonomy database. ’Unclassified’ represents the original scientific name that cannot be classified into any species listed in the GenBank taxonomy database.

This work has largely expanded the number of complete temperate phage genomes. Compared with all the 1,504 GenBank public complete temperate phage genomes of 154 host species and 78 host genera, our result represents an approximately 128-fold (192,326/1,504) increase in the number of complete temperate phage genomes, with an approximately 18-fold (2,717/154) increase in the number of host species, and a 9-fold (710/78) rise in the number of host genera.

By analyzing temperate phage-host interactions (Supplementary Materials, Multiple host infection of temperate phages), we found many temperate phages in common pathogens, such as *Klebsiella pneumoniae*, *Salmonella enterica*, *Enterobacter cloacae*, *E. coli* (Extended Data Table 8). The host species sharing network shows that these temperate phages in common pathogens could also infect other species and genera. In particular, *K. pneumoniae* has the most temperate phages in common with other species in genera *Escherichia*, *Salmonella*, *Enterobacter*, and *Klebsiella*. *E. coli* shares the most temperate phages with *Shigella sonnei* (Figure 3A). The phylogeny network shows that the temperate phages, constituting the largest cluster in the host species sharing network, are also similar at the sequence level (Figure 3B). By referring to the multiple host connection network provided in this study, researchers can use the temperate phages in these common bacteria to infect the uncommon bacteria for a particular research purpose.

**Figure 3.**
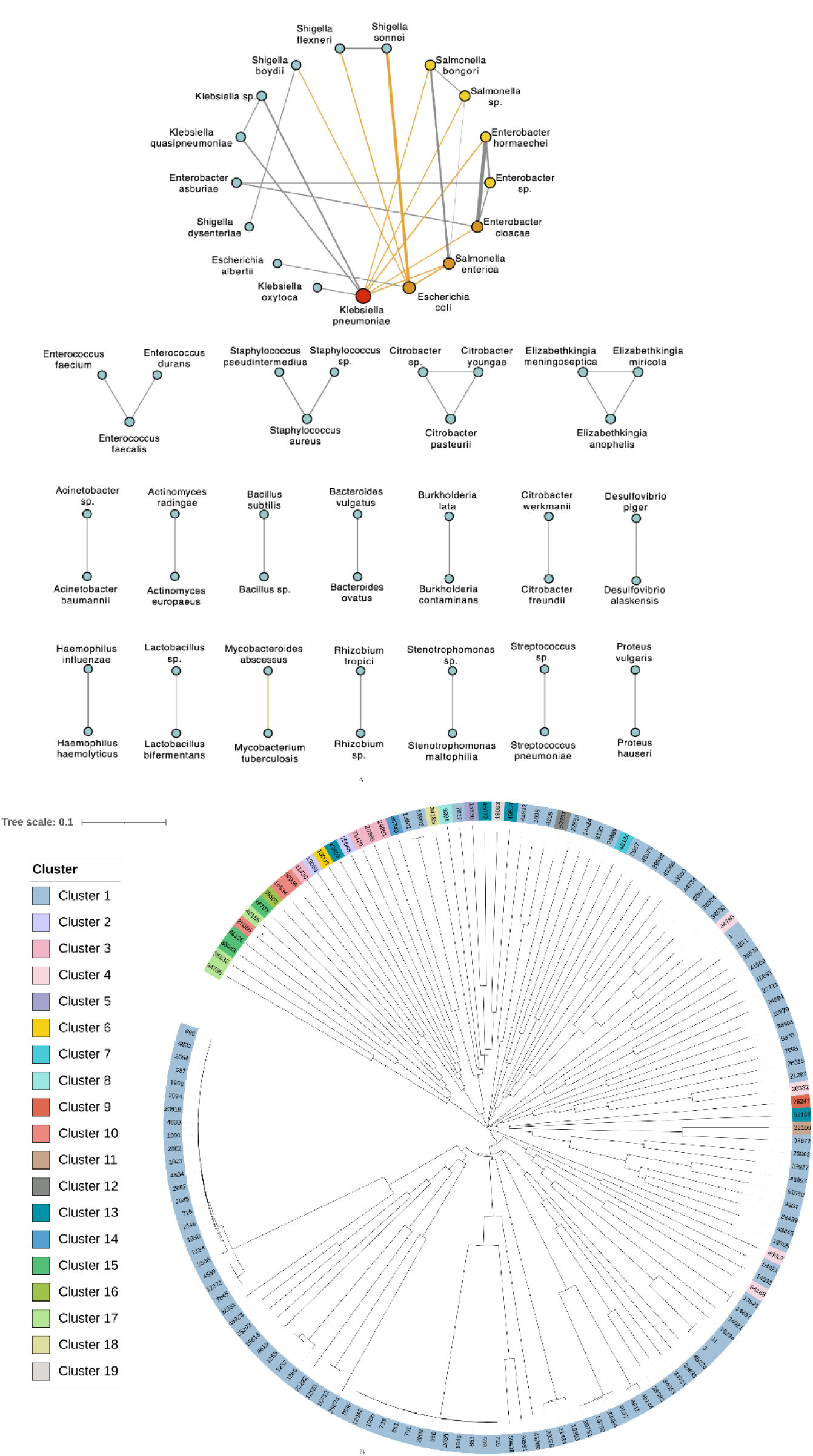
The connected network of temperate phages (edge) with their hosts (nodes). (A) All data labeled as metagenome or unclassified by GenBank are not included. The more temperate phages shared, the thicker the line (the node) is. The gray line represents the same phages identified within the same host genus, while the yellow line represents the phages identified across host genera. (B) Phylogenetic relationships of the phage entries in (A).

Phage-mediated horizontal gene transfer (HGT) plays an essential role in bacterial evolution. This study presented a complex temperate phage-host connection network and antibiotic resistance and virulence gene sharing networks (Supplementary Materials, Horizontal gene transfers mediated by temperate phages), illustrating that temperate phages play essential roles in HGT among host species (Figure 3, Extended Data Figure 7, Extended Data Figure 8). In particular, tetracycline, macrolide, and aminoglycoside resistance genes were shared widely among temperate phages of different host species, as well as the virulence factors of adherence and invasion and secretion systems (Extended Data Figure 4A, B).

Restriction-modification (RM) systems are the best-known bacterial defense mechanisms against foreign DNA, including phages^21^. Usually, the restriction systems consist of a restriction endonuclease responsible for recognizing cleavage sites of a specific DNA sequence. In contrast, the modification systems contain a DNA methyltransferase that modifies unmethylated or hemimethylated DNA within the same sequence ^22,23^. Even though the RM systems are often found on bacterial chromosomes, temperate phages can also protect the bacteria against virulent phage infection by encoding restriction systems ^24^. Given the long and complex dynamic interaction of bacteria and their phages, temperate phages possibly provide a plethora of additional viral defense systems that have yet to be fully identified ^25^. RM systems have been classified into three main classes designated I, II, and III ^26,27^. By aligning with REBASE ^27^, this study preliminarily identified the three RM systems encoded by 2,626 temperate phages, containing 76 host species (Supplementary Materials, Examination of restriction-modification systems encoded by temperate phages). Our research also showed that type II RM systems widely existed in the most temperate phages with different host species (Extended Data Figure 4C). Although RM systems are the weapons for the hosts to fight against phages, our study indicates that temperate phages can protect the hosts with RM systems against other foreign DNA invasions. It is also a self-protection mechanism for the temperate phages to restrict other competing phages.

CRISPR-Cas systems are widely present in bacteria and archaea^28^, constituting a primary and powerful adaptive immune system that helps them escape the threat of viruses and other mobile genetic elements^29^. On the contrary, a remarkable turn of events has been shown that phage also encodes the CRISPR-Cas system to counteract a phage inhibitory chromosomal island of the bacterial host^30^. Six types of CRISPR-Cas systems (I, II, III, IV, V, VI) have been defined and updated based on Cas protein content and arrangements in CRISPR-Cas loci^31^. By searching the CRISPR-Cas systems, our study found 466 temperate phages in 17 host species that encoded both CRISPR array and Cas proteins, constituting ten subtypes within three main types of CRISPR-Cas systems, including types I (I-A∼I-C, I-E∼I-F), II (II-A, II-U), and III (III-A∼III-B, III-U) (Supplementary Materials, Examination of CRISPRs and anti-CRISPRs encoded by temperate phages). Among them, II-U was the most commonly occurring type found in the temperate phages, which infected five host species (Extended Data Figure 4D1, Extended Data Table 11).

On the other hand, anti-CRISPRs provide a powerful defense system that helps phages escape injury from the CRISPR-Cas system^32^. Anti-CRISPR proteins with anti-I-E and anti-I-F activities have been found in *Pseudomonas aeruginosa* phages^33,34^, and those inhibiting type II-C systems have been found in *Neisseria meningitidis*^35^. By searching the anti-CRISPRdb^32^, 1,108 temperate phages in seven host species encode anti-CRISPR proteins inhibiting four important CRISPR systems, including I-E, I-F, II-A, and II-C (Supplementary Materials, Examination of CRISPRs and anti-CRISPRs encoded by temperate phages). The type anti-I-F is the most commonly occurring type found in the temperate phages that infect four host species (Extended Data Figure 4D2, Extended Data Table 12). Also, the temperate phages in *P. aeruginosa* encode anti-CRISPR proteins inhibiting I-E and I-F CRISPR systems, and those in *N. meningitidis* and *Neisseria lactamica* encode anti-CRISPR proteins inhibiting type II-C CRISPR system. Our findings are consistent with the above published results^33–35^. Moreover, it illustrates that not only virulent phages encode the anti-CRISPR proteins, but also temperate phages.

Our findings may facilitate phage-related clinical applications. FMT has been successfully applied to treat CDI patients. However, FMT treatment becomes risky if multidrug-resistant or highly virulent bacteria exist in the transplant’s microbiota. This situation becomes even worse if temperate phages carry important antibiotic resistance and (or) virulence genes. In this study, we found that 34.9% (912/2613) of human gut metagenome samples contained a total of 912 temperate phage genomes (Extended Data Table 7, Extended Data Table 8). The antibiotic resistance (fusidic acid-resistance and macrolide-resistance, Extended Data Table 9) and virulence (adherence and invasion; serum resistance, immune evasion and colonization, Extended Data Table 10) genes in temperate phages may potentially increase the risk of FMT.

Phage therapy is a promising alternative for antibiotics to treat bacterial infections. However, virulent phages for some host bacteria, such as *C. difficile* and *M. tuberculosis*, are difficult to isolate. Therefore, temperate phages against such host bacteria can be converted to the virulent phages to infect those host species. Our study revealed 1,847 temperate phages in *C. difficile* and 1,391 in *M. tuberculosis* (Extended Data Table 7). Their genomes can be further used to transform temperate phages into virulent phages for phage therapy applications.

In this study, we first provide a large temperate phage genome database. Based on the data resource, we then examined the phage genomic diversity, evaluated the diversity of the core region of attachment sites, investigated phage-host interactions, explored mediated HGTs, and examined the encoded RM and CRISPR/anti-CRISPR systems. Our study sheds light on the mosaicity and diversity of temperate phages at the genomic level and illustrates that the phages and their hosts are interconnected through a complex network. Our work paves a way for a better understanding of interactions between temperate phages and their hosts, and further explains the roles that temperate phages may play in bacterial evolvability.

## Methods

### Details of the temperate phage detection method proposed in this study

Our temperate phage detection method includes three main steps (Figure 1).

1. Next-generation sequencing (NGS) data processing. FastQC^1^ and Trimmomatic^2^ are first used to do quality control and NGS reads filtering. The filtered NGS reads are fed into a sequence assembly software, such as Spades, to acquire the assembled scaffolds.
2. Prophage region detection. GLIMMER v 3.02^3^ is used to predict open reading frames (ORFs) on the above bacterial scaffolds. BLASTp is then used to align the ORFs with the local phage protein database which were generated from the GenBank public phage-only protein database with setting the E-value as 1e-10. To acquire as many prophage regions as possible, hypothetical and functional phage genes, such as capsid, terminase, protease, integrase, transposase, lysis, tail, spike, holin, portal, and baseplate, are all taken into consideration. DBSCAN (density-based spatial clustering of applications with noise) is used to identify a phage ORF cluster (Extended Data Table 16). The DBSCAN uses two parameters: the size of the cluster (M) and the radius of the cluster (E). Here, M is defined as the minimum number of phage genes in a possible prophage region and E is the largest distance between any two neighbor genes. A possible prophage region includes more than 30,000 bases before and after the center on the same scaffold when treating a hypothetical phage gene cluster or a functional phage gene cluster as a center. The integrated overlap of any two afore-defined prophage regions by DBSCAN is treated as a rough prophage region. Our method then defines a precise prophage region based on the integration and induction mechanism of a temperate phage (Figure 1). The *attL* and *attR* in Figure 1 are treated as the termini of a temperate phage when the phage inserts into a bacterial chromosome. Considering that *attL* and *attR* have a 14∼50 bp core region, our method defines two sliding windows with the same sizes to detect the core region. The distance between the two sliding windows is defined as d∈[10,000, length (rough prophage region)]. Considering that most prophage genomes are larger than 10,000 bp, the initial value of the distance is set as 10,000 bp. For each d, the sequences in the two sliding windows are compared base by base (Extended Data Figure 10). The comparison stops once the two bases in the two windows are different or reach the end of the window. The repeats with sequence between them are recorded as the ends of the precise prophage region.
3. Temperate phage detection. In our method, a prophage is treated as a temperate phage if the prophage can circulate itself. As shown in Extended Data Figure 11, the 1,000 bp sequences at both ends (A and B) of the precise prophage region are extracted, with their positions being reverted to form a new sequence. Currently, most NGS platforms use paired-end sequencing technology. If this prophage can circulate itself, there must be paired reads that match A and B concurrently, which is the evidence that NGS captures the replication status in the lytic cycle of the temperate phage. Finally, a complete temperate phage sequence is acquired after using an in-house sequence-end-extend script to fill the gap between the paired reads matching A and B.

In our method, the scaffolds with character N inside are processed in two directions (red rectangle in Figure 1). The reason for doing this is that when “N” appears in the scaffold, these Ns cause GLIMMER v 3.02^3^ to predict ORFs differently in the original and reverse orders of the given sequence. That is, we may miss some temperate phages if searching from only one direction. Therefore, reverse complementary forms of these sequences containing character N are taken into consideration so that more temperate phages (within the reverse complementary host genome) are identified. For all the outputs of identical and reverse complementary phage sequences in one host sample (scaffolds with the same SRA number), they were treated as one phage and hence only kept one sequence in our analysis. To balance between quality and variety, a temperate phage sequence is excluded if the number of character N in the sequence takes up more than 5%.

### Biological verification of temperate phages detected in this study

Mitomycin C was used to induce the potential temperate phages in bacteria. The nucleotides in bacterial culture supernatant were extracted to do NGS to identify the temperate phages inside.

Specifically, this biological verification includes five steps:

1. Inducing temperate phages. The monoclonal of bacteria was extracted and put in a 5 ml LB liquid medium. The cultures were incubated at 37°C overnight and then added to 400 ml of fresh LB medium to grow to the stationary phase (OD600=0.5). Mitomycin C was added (1g/ml) into the medium that was then incubated at 37°C for 12 hrs until the medium became clear. The bacterial culture supernatant was then collected.
2. Extracting temperate phages. The above extracted bacterial culture supernatant was then added 23.4g NaCl to acquire a final concentration of 1 mol/l, followed by being stirred and cooled 1 hr in an ice bath. The cooled mixture was then centrifuged at 1,000x g at 4°C for 10 mins. The supernatant in the mixture was then collected and transferred into a clean 500 ml flask. PEG8000 was then added into the supernatant at a ratio of 10mg/100ml, followed by being stirred and cooled 3 hrs in an ice bath. The culture was then centrifuged at 1,000x g at 4°C for 10 mins. After centrifugation, the phage precipitation was then collected and resuspended in SM buffer. The same volume of chloroform was added into the phage precipitation. The phage was then collected at its hydrophilic phase. This phage precipitation was then filtered using a 0.45 ∼m filter and stored at 4°C.
3. Purifying temperate phages. The phage particles were purified by isopycnic centrifugation through CsC1 gradients. The high quality of solid CsC1 was added into the SM buffer and made three different types of CsC1 solutions with different densities (P: 1.45g/ml, P:1.50g/ml, P:1.70g/ml). Each kind of CsC1 solution was added into different Beckman Ultra-Clear centrifuge tubes, which were further added 10 ml phage precipitation prepared at step 2. The tubes were then centrifuged at 25,000 r/min at 4°C for 3 hrs. The phage layer was then sucked up to a new centrifuge tube to remove CsC1 using SM buffer in a 100 kD dialysis bag for 10 hrs. Finally, the pure phage suspension was acquired.
4. Extracting temperate phage genomes. After being added DNase and RNase, the pure phage suspension was then incubated at 37°C for 10 hrs and inactivated at 80°C for 15 mins, followed by extracting the genomes and stored at -20°C.
5. High-throughput sequencing for temperate phage genomes. The preparation of paired-end libraries and whole-genome sequencing by generating 2×150 bp paired-end reads were performed using the Illumina MiSeq sequencing platform by Annoroad Genomics Co., LTD (China).

### Method comparison of temperate phages detected in this study

The bacterial scaffolds assembled in the first step of our temperate phage detection method were used as the input for each prophage prediction method, including Phage_Finder^14^, Prophinder^16^, PHASTER^17^, PhiSpy^3^, VirSorter^18^, Prophage Hunter^4^, and VIBRANT^5^. As Prophage finder^15^ has been out of maintenance, we are unable to run it in our study. CLC Workbench v3 was used to map the NSG reads of the wet-lab induced temperate phages to their host’s assembled scaffolds, with parameter settings length as 1.0 and sequence fraction as 1.0.

### Host assignment based on GenBank taxonomy database

Two .dmp format files named names and nodes in the folder of taxdump were download from https://ftp.ncbi.nlm.nih.gov/pub/taxonomy/. An in-house script in Python 3.6 was used to assign the host for each phage genome. All host ranks of each phage are listed in Extended Data Table 7.

### Analysis of temperate phage genomes

All the primary analysis in this study, including the distributions of length and GC content, as well as genome classification, was conducted using the Python programming language (version 3.6.0) implemented through Geany 1.33, with the related images being generated using the ggplot2 package^5^. Genome annotations were conducted using online annotation server RAST^6–8^, while the genomic mapping was generated using an in-house Perl script. The connected network was also implemented by an in-house developed script in Python 3.6 and used to display the connections between phages and their multi-hosts, where nodes represent hosts while edges (lines) are phages. A cluster is formed as long as the hosts inside can be infected by at least one same temperate phage. Multiple alignments of the temperate phage sequences were implemented by MAFFT v5^9^. The phylogenetic relationship was analyzed by the neighbor-joining method implemented in MEGA X^10^ with default settings and then displayed using iToL^11^.

### Detection of antibiotic resistance genes, virulence genes, genomic islands, restriction-modification, CRISPR and anti-CRISPR systems

Antibiotic resistance genes were identified using the database in ResFinder^12^, and virulence gene identification was conducted by VFDB^13^. IslandPath-DIMOB^14^ was used to detect genomic islands with the input of phage genomes annotated by Prokka^15^. The restriction-modification (RM) systems were searched against REBASE^16^. BLAST was then used to align all the temperate phage sequences with the above databases by setting e-value as 1e-5 and choosing 90% as lower bound for sequence coverage. Moreover, the restriction (R) and the modification (M) enzyme genes are often tightly linked to form a restriction-modification (RM) gene complex^17^. The type I systems also have sequence recognition (S) subunit genes to form multi-subunit enzymes for modification (SM) or restriction (SMR)^18,19^. These genes are found to be separated by an intergenic region, such as 56 bp^20^, 76 bp^21^, 109 bp^22^, 110 bp^23^, 330 bp^24^, 365 bp^25^. Therefore, we set the upper bound of the distance between such genes as 400 bp. The CRISPR-Cas systems were searched using CRISPRCasFinder^26^. Only the temperate phages containing both the CRISPR array and the Cas proteins are counted in our study. The anti-CRISPR systems were searched against the database Anti-CRISPRdb^27^ by BLAST to align all the temperate phage sequences with setting e-value as 1e-5 and choosing 90% as lower bound for sequence coverage.

### Data availability

The source code implementing our method can be downloaded from https://github.com/NancyZxll/temperate-phage-active-prophage-detection. All the raw data for method validation and output of each tools have been made available at https://phage.deepomics.org/files/. All the temperate phage genome sequences detected in our study are freely available via https://phage.deepomics.org/. All additional in-house scripts and other relevant data are available from the corresponding author on request.

## Acknowledgement

This work was provided by the National Key Research and Development Program of China (Grant No. 2017YFF0108605 and 2018YFC160025, 2018YFA0903000) and the National Natural Science Foundation of China (Grants 31900489, 81672001 and 81621005). The work of X. J. was supported by the Intramural Research Program of the National Library of Medicine, National Institutes of Health.

## Author contributions

X.Z., R.W. designed and performed bioinformatics analyses. X.J. collected all datasets. M.W. curated data for the study. X.X. wrote the essential phage detection program, X.Z., Q.S. and Q.N. contributed to revise the program. X.Z., S.Z., S.T. and F.W. contributed to test the program. X.F. developed the website. Y.H. conducted the wet-lab experiments. X.Z. wrote the manuscript. Y.T, J.W., A.M.K., S.L., Y.C., and X.J. revised the manuscript. Y.T., S.L., and S.P. conceived and supervised the project. All authors approved the final manuscript.

## Competing interest

The authors declare no competing interests. Correspondence and requests for materials should be addressed to Y.T.

## Supplementary information

### Verification of our temperate phage detection method

Our method detected 17 temperate phages from 15 out of the total 148 bacteria preserved in our laboratory, with two temperate phages induced from bacterium No.1367 and bacterium No.1369, separately (Extended Data Table 1 and Extended Data Figure 1). To verify our results, we then induced the temperate phages in the 15 bacteria by wet-lab experiments (Materials and Methods). The extracted temperate phages were then sequenced using the MiSeq next-generation sequencing (NGS) platform. The NGS reads of wet-lab induced temperate phages were mapped to their hosts’ assembled scaffolds. The mapped positions of the NGS reads are converted into corresponding temperate phage sequence positions, which are treated as ground truth (Extended Data Figure 1, labeled as “Biological experiment”). Compared with Prophage finder^15^, Prophinder^16^, PHASTER^17^, PhiSpy^3^, VirSorter^18^, Prophage Hunter^4^, and VIBRANT^5^, our method identified the same sequence boundaries as the wet-lab biological experiments (Extended Data Figure 1 and Extended Data Table 2). This result illustrates that our method can acquire accurate complete temperate phage genomes. We also compared the calculation time and memory usage of the offline tools, including VirSorter, VIBRANT, PhiSpy, Phage_finder, and our method (Extended Data Figure 2, details in Extended Data Table 3). Due to the consideration of sequence boundaries (Table 1), VirSorter and our method take more calculation time than other three tools. Notably, our method takes much less time than VirSorter and comparable to VIBRANT that does not consider sequence boundaries. Because our method needs bacterial NGS reads to detect sequence circulation, it needs the most memory space (∼700MB) among all the offline tools.

### Expansion of temperate phages

Using our temperate phage detection method, we identified 192,326 complete temperate phage sequences within 2,717 host species. By grouping the same/reverse complementary sequences as one entry, there are 66,823 nonredundant complete temperate phage genomes in total. Compared with the 1,504 GenBank public temperate phage genomes having 154 host species, this represents an approximately 128-fold increase in the number of complete temperate phage sequences with an 18-fold increase in the number of host species.

### Temperate phages extracted from different hosts

As phages isolated in nature rarely have near-identical genomes, the term “species” is not frequently used when describing relationships among phages^2^. Here we use the taxonomic ranks of the hosts to illustrate their relationships. For the 192,326 complete temperate phage sequences, at the level of species, the temperate phages identified in *Salmonella enterica* represents 72% (138,316/192,326) of the total phages (Extended Data Table 7), rendering that *S. enterica* contains the most temperate phages in public genomic database. It may be tempting to regard the reason that *S. enterica* has the most host strains, by taking up 34.3% (248,957/726,900) of all host strains and 2.81x (248,948/88,468) as many as the top second host species (Extended Data Figure 3A, Extended Data Table 8).

For the total number of temperate phages identified in their respective host groups (Extended Data Figure 3B left panel in each taxonomic rank, Extended Data Table 8), at the level of species, 55.5% *S. enterica*, along with its subspecies, contain 138,316 complete temperate phages, followed by *K. pneumoniae* (40.5%, 8,471) and *E. coli* (9.4%, 8,340). At the level of genus, *Salmonella*, *Klebsiella*, and *Escherichia* reach the top three: 138,582 temperate phages were identified from 55.4% *Salmonella*, 9,085 temperate phages were identified from 40.6% *Klebsiella*, and 8,362 temperate phages were identified from 9.4% *Escherichia*. At the level of family, *Enterobacteriacea* (159,346 phages identified from 374,762 host strains), family *Listeriaceae* (5,381 phages from 30,090 host strains), and family *Staphylococcaceae* (4,444 phages from 60,236 host strains) are the top three hosts containing the most bacteria. At the level of order, *Enterobacterales* (160,416 phages from 378,602 host strains), order *Bacillales* (14,163 phages from 92,856 host strains), and order *Lactobacillales* (3,406 phages from 46,249 host strains) reach the top three; class, *Gammaproteobacteria* (164,655 phages from 408,466 host strains), class *Bacilli* (14,163 phages from 139,105 host strains), and class *Actinobacteria* (2,579 phages from 56,440 host strains) reach the top three; phylum, *Proteobacteria* (169,490 phages from 503,942 host strains), phylum *Firmicutes* (16,673 phages from 152,801 host strains), and phylum *Actinobacteria* (2,608 phages from 56,482 host strains) reach the top three.

However, for the average number of temperate phages identified per host strain, many rare host types top the list at each rank-level (Extended Data Figure 3B right panel in each taxonomic rank). For example, at the species level, four temperate phages were identified in each of the strains in *Pseudoalteromonas sp*. P1-8, *E. cloacae complex* sp. ECNIH11, *Pseudomonas* sp. SJZ074, etc., respectively. At the genus level, *Nautella*, *Parvimonas*, *Niabella* reach the top with three temperate phages being identified in each of their strains. At the family level, in each strain of *Marinifilaceae*, *Jiangellaceae*, *Tissierellaceae*, *Thiolinaceae*, and *Chthoniobacteraceae* identified two temperate phages; thus, these families reach the top. At the order level, two temperate phages were identified in each strain of *Chthoniobacterales* and *Jiangellales*, which reach the top. At the class level, *Spartobacteria* reaches the top with two temperate phages being identified in the only strain (*Chthoniobacter flavus*) in this class. At the phylum level, 1.5 temperate phages were identified in each strain of *Synergistetes*, to make this phylum reach the top. These results signify that even though a host group is integrated by the most temperate phages in total, it does not necessarily mean that each host strain in this group is integrated by the most temperate phages.

### Temperate phage genome entries

For simplicity and clarity, phage entry will be used as a short form for nonredundant complete temperate phage. For the 66,823 phage entries, based on the GenBank bacterial taxonomy at the time, except the hosts from metagenome or named as bacterium by data submitter, all hosts are divided into 34 phyla, 51 classes, 229 families, 112 orders, 708 genera, 2,717 species (Extended Data Table 6). As shown in Extended Data Figure 3C, about 76.7% of all the temperate phage entries infect the host phylum of *Proteobacteria*; while the host phylum of *Deferribacteres* contains the most abundant temperate phages per bacterium (∼1.33 phages infect one bacterium). Notably, at the level of species, *S. enterica* along with its subspecies harbour 32,520 temperate phage entries with each strain carrying 0.131 phage entries on average, followed by *E. coli* (infected by 6,042 temperate phage entries, 0.068 entries per strain on average) and *Listeria monocytogenes* (infected by 3,365 temperate phage entries, 0.151 entries per strain on average). While *K. pneumoniae* contains the second most temperate phages (8,470) mentioned in the above section of “Temperate phages extracted from different hosts”, this species includes 3,169 phage entries with 0.151 entries per strain, listed in the fourth place.

In conclusion, *S. enterica*, *K. pneumoniae*, and *E. coli* are the top three species that contain the most temperate phages. In contrast, *S. enterica*, *E. coli*, and *L. monocytogenes* are the top three species that include the most abundant temperate phages (the most temperate phage entries). However, unlike the statistics of the total number of phages, many rare types of bacteria reach the top list when calculating the number of infected temperate phages/phage entries per host strain on average.

### Ranges in genome size and GC content of temperate phage entries

Temperate phage entries have a wide range of genome sizes, especially those associated with *L. monocytogenes*, from 9,451 bp to 99,951 bp (Extended Data Table 6, Figure 3A). The smallest temperate phage genome is that from *Campylobacter*, with 9,316 bp. At the other end, *S. enterica* has the largest temperate phages, with some at 99,989 bp. In the aspect of GC content, the temperate phage infecting the species *Brachyspira murdochii* has the lowest GC content of 23.89%, while that infecting *Clavibacter michiganensis* has the highest GC content of 78.52% (Extended Data Table 6). Within the top ten most hosts, the human gut metagenome contains the broadest GC content range of temperate phages (Figure 3B), which is consistent with the fact that the metagenome contains various bacteria. On the contrary, the temperate phage genomes in the top three hosts, including *S. enterica*, *E. coli*, and *L. monocytogenes* have a relatively constant GC content.

### Multiple host infection of temperate phages

Of the 66,823 nonredundant complete temperate genomes (temperate phage entries), 252 have multiple hosts (Extended Data Table 6). The genome sizes of these 252 temperate phage entries range from 9,430 bp to 82,029 bp, while the phages in multiple hosts of *S. sonnei* and *E. coli* have the broadest range from 10,245 bp to 82,021 bp (Extended Data Table 6). The GC contents of the 252 temperate phage entries range from 26.0% to 69.9%, while the phages in multiple hosts of *S. enterica* and *Salmonella sp.* have the widest range from 40.3% to 51.0% (Extended Data Table 6). Interestingly, 15 temperate phage entries identified in human gut metagenome were also found in *E. coli* (7 temperate phages), *E. cloacae* (3 temperate phages), *Enterobacter hormaechei* (2 temperate phages), *Enterobacter sp*. (1 temperate phage), *Pantoea agglomerans* (1 temperate phage), *K. pneumoniae* (1 temperate phage), *Citrobacter koseri* (1 temperate phage), and *Staphylococcus aureus* (1 temperate phage) (Extended Data Table 6).

After removing the phage entries whose genera contain “NA”, “metagenome”, and “uncultured bacterium”, 196 of the total 252 phage entries were considered in phylogenetic analysis. The phylogenetic relationship shows that 26.5% (52/196) of the 196 phage entries found in the genus of *Salmonella* share high sequence similarity (Extended Data Figure 5A). Moreover, the phage entries with cross-genera hosts were found in the genera of *Klebsiella*, *Salmonella*, *Escherichia*, *Shigella*, *Enterobacter*, *Mycobacterium*, and *Mycobacteroides*; while most them infecting *Escherichia* also tends to infect *Shigella*. (Extended Data Figure 5B).

A network was built to display cross-species connection, where nodes (drawn as points) representing hosts and edges (connections between nodes, drawn as lines) corresponding to the temperate phage entry (Figure 3). In total, 147 of 196 temperate phage entries consist of 19 cross-species clusters (Figure 3). The temperate phages in the biggest cluster share the most hosts in different species (Figure 3, Extended Data Table 6). For example, the temperate phages in *K. pneumoniae* appear closely related to *S. enterica*, *S. sp.*, *Salmonella bongori*, *Klebsiella quasipneumoniae*, *Klebsiella sp*., *Klebsiella oxytoca*, *Enterobacter hormaechei*, *E. cloacae*; those in *E. coli* also infect *S. sonnei*, *Shigella flexneri*, *Shigella boydii, S. enterica*, and *Escherichia albertii*; the temperate phages in *S. enterica* can infect *K. pneumoniae*, *E. coli*, *S. bongori*, and *S. sp*.

### Examination of core regions of phage-chromosomal attachment sites

Temperate phage can lysogenize bacterial strains by a site-specific recombination process ^3,4^. The specific site is called phage-chromosomal junction fragments/ attachment site; the repeated sequence in both temperate phage and bacterium is called the core region of an attachment site (Figure 1).

The sizes of the core regions range from 13 bp to 248 bp (Extended Data Table 13). Within the top 20 most host species, the core regions of temperate phages in *S. enterica* and *E. coli* have the broadest size range, followed by *S. sonnei*, *L. monocytogenes*, *Haemophilus influenzae* (Extended Data Figure 6A). On the other hand, the core regions of temperate phages in *M. tuberculosis* and human gut metagenome have the widest GC content range (Extended Data Figure 6B). It is noted that the core regions of most temperate phages identified in marine metagenome have a relatively stable size of 77 bp but a wide range of GC content (Extended Data Figure 6, Extended Data Table 13).

### Horizontal gene transfers mediated by temperate phages

The process in which a temperate phage inserts itself into host bacteria and becomes a prophage is called phage-lysogenic conversion^5^. During the phage-lysogenic conversion, the temperate phage plays a major role in the horizontal gene transfer (HGT)^6,7^, which is a process transforming benign bacteria into pathogens by transferring antibiotic resistance genes ^8,9^ and introducing virulence factors^10,11^. On the other hand, genomic islands, the syntenic gene blocks formed by HGT, also significantly impact the genome plasticity and evolution, and disseminate antibiotic resistance genes and virulence factors^12^. Temperate phages are proven to contain genomic islands^13,14^. By considering the impact of HGT factors mediated by temperate phages on bacterial evolution and survival, this study analyzed the basic characteristics of functional genes, including antibiotic resistance genes and virulence factors, as well as genomic islands.

### Antibiotic resistance genes in temperate phages

The antibiotic resistance genes in the 11 antibiotic resistance types listed in ResFinder^15^ were identified in 546 temperate phages of 42 host species. The temperate phages in *Escherichia coli* and *Salmonella enterica* contain the most antibiotic resistance gene types (eight gene types for each). Specifically, the eight gene types in *E. coli* were identified in 66 temperate phages, with each containing 1∼5 gene types (Extended Data Figure 4A, Extended Data Table 9). For *S. enterica*, the eight gene types were identified in 53 temperate phages, with each temperate phage containing 1∼4 gene types. Seven gene types were identified in 54 temperate phages in *S. sonnei*, with each phage having 1∼4 gene types (Extended Data Figure 4A, Extended Data Table 9). On the other hand, there are three antibiotic resistance gene types widely identified in the temperate phages within different host species, including tetracycline resistance genes identified in 97 temperate phages within 19 host species, macrolide resistance genes identified in 36 phages within 15 host species, and aminoglycoside resistance genes identified in 192 phages within 12 host species. It illustrates that these kinds of antibiotic resistance genes may play important roles in the co-survival of bacteria and their prophages when facing antibiotic. By contrast, the colistin resistance and the fosfomycin resistance genes were only identified in one temperate phage of *E. coli* and *B. anthracis*, respectively.

Moreover, all temperate phages can be classified into three gene-sharing groups (Extended Data Figure 7). In the connected network, there is an edge connecting the two host species if their temperate phages contain the same antibiotic resistance gene. The temperate phages in *Staphylococcus pseudintermedius*, *Staphylococcus epidermidis* and *S. aureus*, which are the three species in the genus *Staphylococcus*, share the most antibiotic resistance genes (the gray triangle in group I). In group II, the temperate phages in *E. coli*, *S. enterica*, *S. flexneri*, and *S. sonnei* share the most antibiotic resistance genes; while the temperate phages in *Klebsiella aerogenes* and *Shigella dysenteriae* tend to share many genes with the above four species, respectively. The temperate phages in *Bacillus* share genes only with the others in the same genus (group III).

Notably, the temperate phages in *C. difficile* contain macrolide, tetracycline, and aminoglycoside resistance genes (Extended Data Figure 4A, Extended Data Table 9). Those temperate phages may increase the types and abilities of antibiotic resistance in *C. difficile* strains. Moreover, the temperate phages in *Actinomyces europaeus*, *Citrobacter freundii*, *Campylobacter* sp., *Campylobacter jejuni*, *Enterococcus hiare*, *Streptococcus agalactiae*, *Streptococcus suis*, and *Streptococcus mitis* share the same antibiotic resistance genes with those in *C. difficile*. Therefore, HGT may more tend to happen between those species and *C. difficile*. The above findings should be taken into consideration when treating *C.difficile* infection (CDI) patients, such as using fecal microbiota transplantation (FMT).

### Virulence genes in temperate phages

The virulence genes of the 24 virulence factors listed in VFDB^16^ were identified in 6,403 temperate phages infecting 123 host species in total. Within the top 50 most host species, nearly 50% (11/24) virulence factors were identified in 3,606 temperate phages of *S. enterica* with each phage containing 1∼4 virulence factors. The temperate phages infecting *S. enterica* take up more than 50% of the total number of temperate phages which have virulence genes (Extended Data Table 10). *E. coli* reaches the second place with nine virulence factors identified in its 168 temperate phages, followed by *S. aureus* and *K. pneumoniae* (eight factors identified in 1,380 temperate phages, and eight factors in 19 phages, respectively) (Extended Data Table 10). On the other hand, the temperate phages infecting the most host species have the virulence factors of adherence and invasion (531 phages within 88 host species), followed by factors of secretion system (2,389 phages within 27 host species); chemotaxis, motility, export apparatus and enhance binding to erythrocytes (36 phages in 21 host species); and serum resistance, immune evasion and colonization (844 phages in 19 host species) (Extended Data Figure 4B, Extended Data Table 10). By contrast, the virulence factors of cell surface components, LEE-encoded TTSS effectors, and superantigens were only found in the temperate phages of *Streptococcus pyogenes*; the virulence factors of Non-LEE encoded TTSS effectors, Autotransporter, and Protease, were only found in 15, six, and one phages of *Escherichia coli*, separately; the factors of surface protein anchoring and superantigen were only identified in two phages of *S. aureus* and *Streptococcus pyogenes*, individually; the factors of secreted proteins and antimicrobial activity were only found in one phage from *M. tuberculosis* and *P. aeruginosa*, respectively (Extended Data Figure 4B, Extended Data Table 10). Within the top 50 most host species, the virulence factor of toxin was found in temperate phages of nine host species, including those of *S. aureus* (1204 phages), *S. enterica* (109 phages), *E. coli* (91 phages), *Corynebacterium diphtheriae* (38 phages), *Acinetobacter pittii* (13 phages), *Corynebacterium ulcerans* (13 phages), *K. pneumoniae* (2 phages), *Aeromonas hydrophila* (1 phage), and *Corynebacterium xerosis* (1 phage) (Extended Data Figure 4B, Extended Data Table 10). Moreover, the temperate phages in *B. anthracis* contain the virulence factor of serum resistance, immune evasion (Extended Data Table 10). Compared with the antibiotic resistance genes contained in the temperate phages of *C. difficile*, the temperate phages in this host species only include the virulence factor of Toxin (Extended Data Table 10).

In the aspect of virulence gene sharing, all the temperate phages constitute 11 groups in total. The most temperate phages share genes in group I: the temperate phages from *Mycobacterium gordonae* share the most virulence genes with the phages infecting other host species (Extended Data Figure 8). Interestingly, the temperate phages within two species of the genus *Aeromonas*, within three species of the genus *Corynebacterium*, within three species of the genus *Neisseria*, and within four species of the genus *Pseudomonas*, share genes in their respective same host genus (Extended Data Figure 8). Even though the most virulence factors (11 out of total 24 factors) were identified in the most temperate phages targeting the species of *Salmonella enterica* (3,950 out of all the 6,403 phages), these phages only have few virulence genes shared with other species. To be specific, these genes were shared with the phages from *S. sp*., *K. pneumoniae*, *Cronobacter sakazakli*, *Enterobacter reggenkampil*, *E. cloacae*, *E. coli*, *S. flexneri* (Extended Data Figure 8, left in middle panel).

### Genomic islands encoded in temperate phages

In total, 10,178 GIs were found in all temperate phages. All the GIs from different phages were grouped into one entry if their sequences were the same or reverse complementary. The 10,178 GIs were grouped into 4,223 GI entries, which were acquired from the temperate phages of 229 host species, belonging to 102 genus and 65 families (Extended Data Table 14). The GI entries identified from phages of *S. enterica* take up half of the total number (55.8%, 2,355/4,223), followed by *E. coli* (18.5%, 781/4,223) and *L. monocytogenes* (3.9%, 166/4,223). For the size distributions of the top 20 host species containing the most GI entries, these in phages of the species *S. sonnei* have the widest size range, while with a relatively consistent GC content (Extended Data Figure 9A, B). Even though the phages in *S. enterica* contain the most GI entries, most of them have a relatively narrow size range with GC content from 34.7% to 62.2% (Extended Data Figure 9A, B). Interestingly, the GI entries identified in temperate phages from human gut metagenome have a relatively narrow size range, from 2,750 bp to 9,933 bp, while with the widest GC content range, from 25.8% to 61.2% (Extended Data Figure 9A, B). Like the temperate phages from the human gut metagenome, the GI entries in the temperate phages of *M. tuberculosis* have a relatively narrow size range while a wide GC content range (Extended Data Figure 9A, B).

Even though there are 4,223 GI entries in total, only 0.7% (31/4223) were shared among phages in different host species and constitute six groups (Extended Data Figure 9C). In particular, the temperate phages from *Bacteroides vulgatus* share the most GI entries with those from the other five species from the same genus of *Bacteroides*, and two species from the genus of *Parabacteroides*. Despite the most GI entries identified in the temperate phages from *Salmonella enterica* (2,355 out of total 4,223 GIs), only 0.3% of them (8/2,355) were shared with species *S. sp*., *S. bongori*, and *L. monocytogenes* (Extended Data Figure 9C).

### Examination of restriction-modification systems encoded by temperate phages

This study preliminarily identifies RM systems encoded by all the 66,823 temperate phage entries. Basically, RM systems have been classified into three main classes designated I, II, and III^17,18^. By aligning with REBASE^18^, the three RM systems have been identified in 2,626 temperate phages in 76 host species. Particularly, within the top 50 host species where temperate phages contain the most RM systems, Type II RM systems were most widely identified in 1,967 temperate phages, by taking up 82% (41/50) of total top 50 host species; while type I RM system identified in 56 phages taking up 18% (9/50) of total top 50 host species; type III identified in 577 phages taking up 22% (11/50) of total top 50 host species (Extended Data Figure 4C, Extended Data Table 15). In addition, all the three RM systems have been identified in the temperate phages in *S. enterica* (type I RM system identified in 11 phages, type II RM system identified in 631 phages, type III RM system identified in 33 phages), *E. coli* (type I in 36 phages, type II in 131 phages, type III in 65 phages), *K. pneumoniae* (type I in one phage, type II in 12 phages, type III in 2 phages) (Extended Data Figure 4C, Extended Data Table 15). Two RM systems were identified in five species, including *L. monocytogenes* (type II in 921 phages, type III in 466 phages), *E. albertii* (type II in one phage, type III in one phage), *Ilyobacter polytropus* (type I in one phage, type III in one phage), *Listeria innocua* (type II in one phage, type III in one phage), and *Paracoccus sanguinis* (type II in one phage, type III in one phage) (Extended Data Figure 4C, Extended Data Table 15).

### Examination of CRISPRs and anti-CRISPRs encoded by temperate phages

In our study, three types of CRISPR-Cas systems, including five, two, and three subtypes in type I, II, and III, respectively, have been identified in 466 temperate phages (Fig 10A, Tab. S11). The temperate phages in *S. enterica* encode the most CRISPR-Cas system with type III-A, followed by the phages in *Campylobacter fetus* encoding type I-B system. Type II-U was found in the temperate phages with the most host species (five in total), followed by type I-E found in the phages with four host species (Extended Data Figure 4D1, Extended Data Table 11). Most CRISPR-Cas systems were only identified in the temperate phages infecting one or two host species (Extended Data Figure 4D1, Extended Data Table 11). Type I-A was only identified in the temperate phages of *S. enterica*; type I-F was only in the phages of *Vibrio cholerae*; type III-U was only in the phages of *Campylobacter lari* (Fig 10A, Tab. S11).

On the other hand, anti-CRISPRs provide a powerful defense system that helps phages escape injury from the CRISPR-Cas system^19^. Anti-CRISPR proteins with anti-I-E and anti-I-F activities have been found in *P. aeruginosa* phages^20,21^; those inhibiting the type II-C system have been found in *N. meningitidis*^22^. By searching the anti-CRISPRdb^19^, 1,108 temperate phages in seven host species were found to encode anti-CRISPR proteins (Extended Data Figure 4D2, Extended Data Table 12). Specifically, the temperate phages in *P. aeruginosa* encode anti-CRISPR proteins inhibiting I-E and I-F CRISPR systems; those in *N. meningitidis* and *N. lactamica* encode anti-CRISPR proteins inhibiting type II-C CRISPR system. These findings are consistent with the above published results. It illustrates that not only virulent phages, but also the temperate phages in those species encode the related anti-CRISPR proteins. Moreover, the temperate phages in *Acinetobacter radioresistens* were found to encode anti-CRISPR proteins with anti-I-F activity, those in *Klebsiella penumoniae* encode anti-CRISPR proteins inhibiting type I-E and I-F systems. The temperate phages in *L. monocytogenes* encode anti-CRISPR proteins inhibiting type II-A system, and those in *Pasteurella multocida* encode anti-CRISPR proteins with anti-I-F activity.

## Supplementary tables

Extended Data Table 1. Laboratory-preserved bacterial strains used in the biological verification.

Extended Data Table 2. The prophage identified in each bacterial chromosome by different methods. Pos. is short for position.

Extended Data Table 3. Calculation time and memory requirement of the offline tools. The system used in this comparison is Linux Ubuntu 14.04.6 LTS, 755G RAM, 80x 2.0GHz Intel Xeon E7-4820 20-core. The number in the table header represents the bacterium NO. PHASTER, Prophinder, Prophage Hunter, and Prophage finder (currently out of maintenance) are web-based tools, therefore they are not considered in the comparison.

Extended Data Table 4. GenBank accession number and depiction of known temperate phage sequences. These sequences were acquired by searching ((((bacteriophage) AND integrase AND (complete genome)) AND Viruses[Organism]) NOT RefSeq) on GenBank.

Extended Data Table 5. The criteria to choose raw NGS data on GenBank.

Extended Data Table 6. Statistics of temperate phage entries, including sequence lengths and GC contents, and taxonomic classifications for their host scientific names. The scientific name is from the column extracted from GenBank-downloaded runinfo.csv. NA, not applicable. \, phage was extracted from eukaryote associated bacteria. Ignoring the hosts that contain “NA” and “meta data”, 240 phage entries have multiple host species, including (uncultured) bacterium and gut metagenome. All sp. species in the same genus are summed up in one species labeled as species sp. in this genus.

Extended Data Table 7. Host assignment of all the 192,326 temperate phages. NA, not applicable. Peptoclostridium difficile is a synonym for C. difficile. They both belong to the genus Clostridioides (76). Star represents that the host of the phage’s host is not consistent with the original host’s name when the data was submitted to GenBank.

Extended Data Table 8. Group of phages and hosts by host scientific name. The scientific name is from the column extracted from GenBank-downloaded runinfo.csvs. NA, not applicable.

Extended Data Table 9. Antibiotic resistance genes carried by temperate phages. Each gene was classified into its antibiotic resistance type listed in ResFinder (53). NA, not applicable. \, phage was extracted from eukaryote associated bacteria.

Extended Data Table 10. Virulence genes carried by temperate phages. Each gene was classified into the related virulence factor listed in VFDB (57). NA, not applicable. \, phage was extracted from eukaryote associated bacteria.

Extended Data Table 11. CRISPR-Cas systems carried by temperate phages.

Extended Data Table 12. Anti-CRISPR proteins carried by temperate phages.

Extended Data Table 13. Core region of attachment sites of temperate phages, including sequence lengths and GC contents, and taxonomic classifications for their host scientific names. The scientific name is extracted from GenBank-downloaded runinfo.csv. NA, not applicable. The ’\’ means that phage was extracted from eukaryote associated bacteria.

Extended Data Table 14. Genomic islands carried by temperate phages, including sequence lengths and GC contents, and taxonomic classifications for their host scientific names. The scientific name is the column extracted from GenBank-downloaded runinfo.csv. NA, not applicable. The ‘\’ means that phage was extracted from eukaryote associated bacteria.

Extended Data Table 15. Restriction and modification (RM) systems carried by temperate phages. The genes coding for restriction endonuclease (R) and methyltransferase (M), their classifications, and host taxonomic classification of each temperate phage having the gene(s). NA, not applicable. \, phage was extracted from eukaryote associated bacteria.

Extended Data Table 16. DBSCAN algorithm pseudo-code. W represents phage genes, E is the radius of cluster, M is the total number of phage genes in one cluster, R is cluster.

## Supplementary figures

**Extended Data Figure 1.**
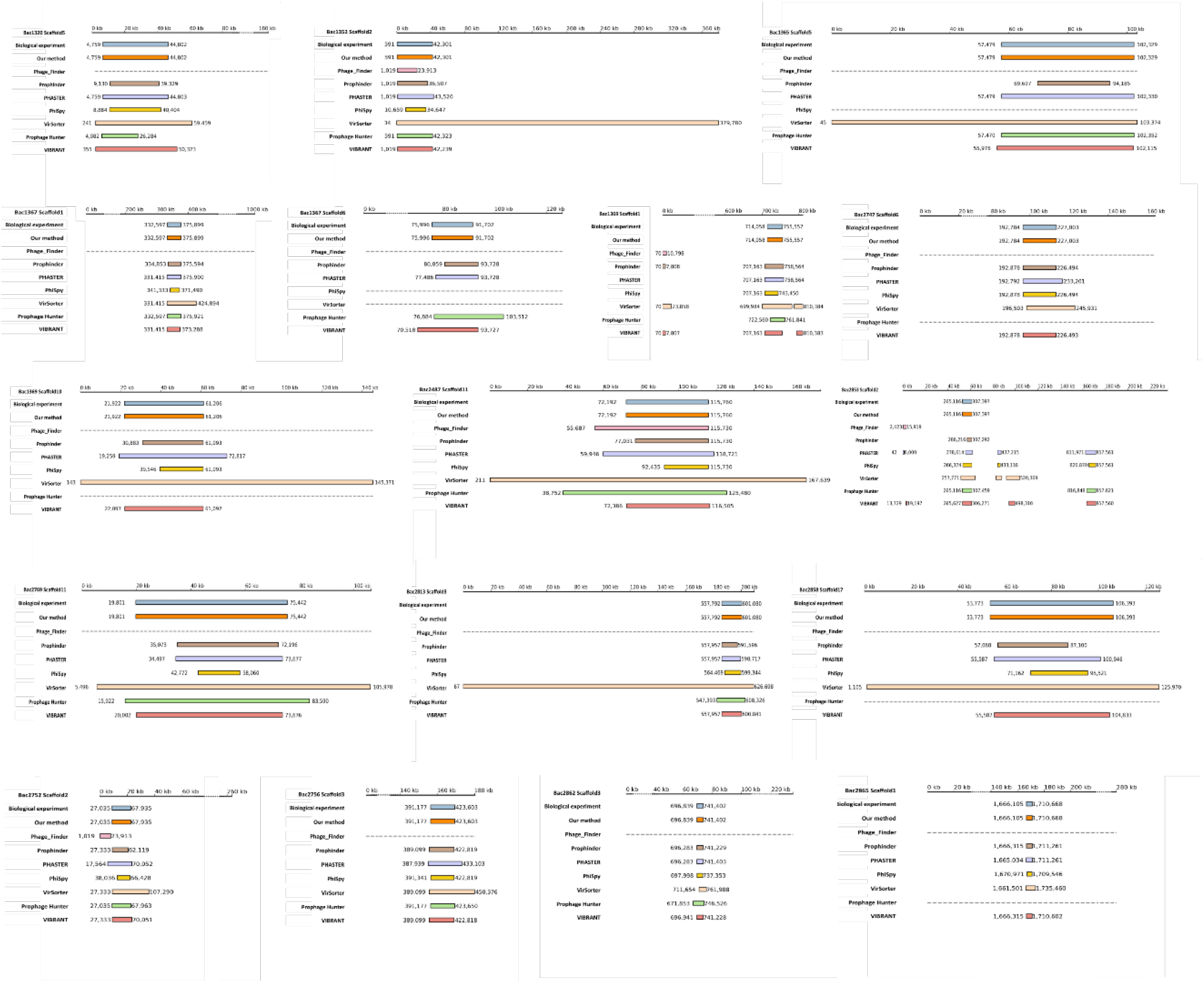
Temperate phage identification by different methods. The scaffold NO. of the bacterium NO. is named as ‘BacNo.ScaffoldNo.’. The rectangle represents the prophage sequence identified by each of the tools. The start and end positions are labeled near the ends of the sequence.

**Extended Data Figure 2.**
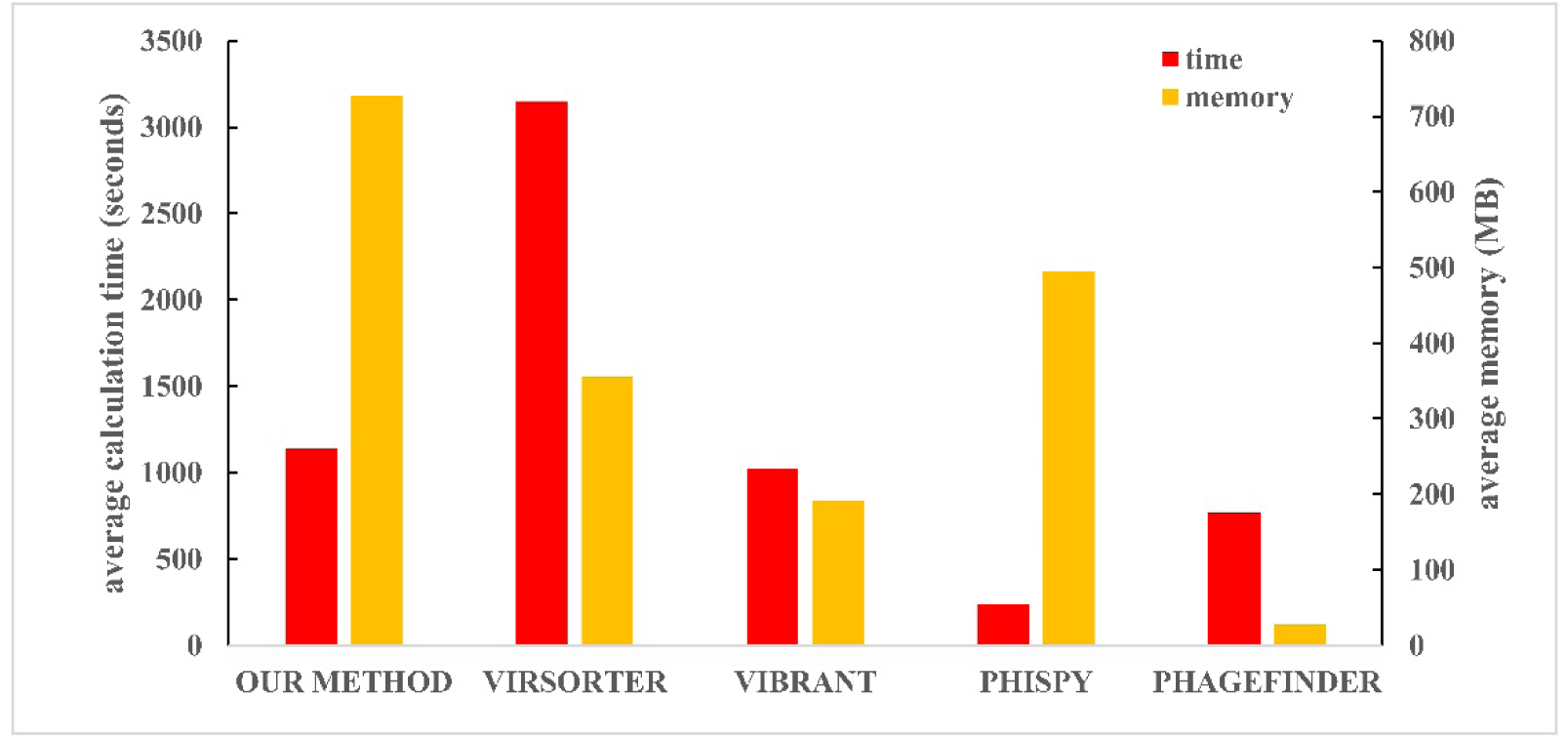
Average calculation time (seconds) and memory usage (MB) of our method and every offline tool.

**Extended Data Figure 3.**
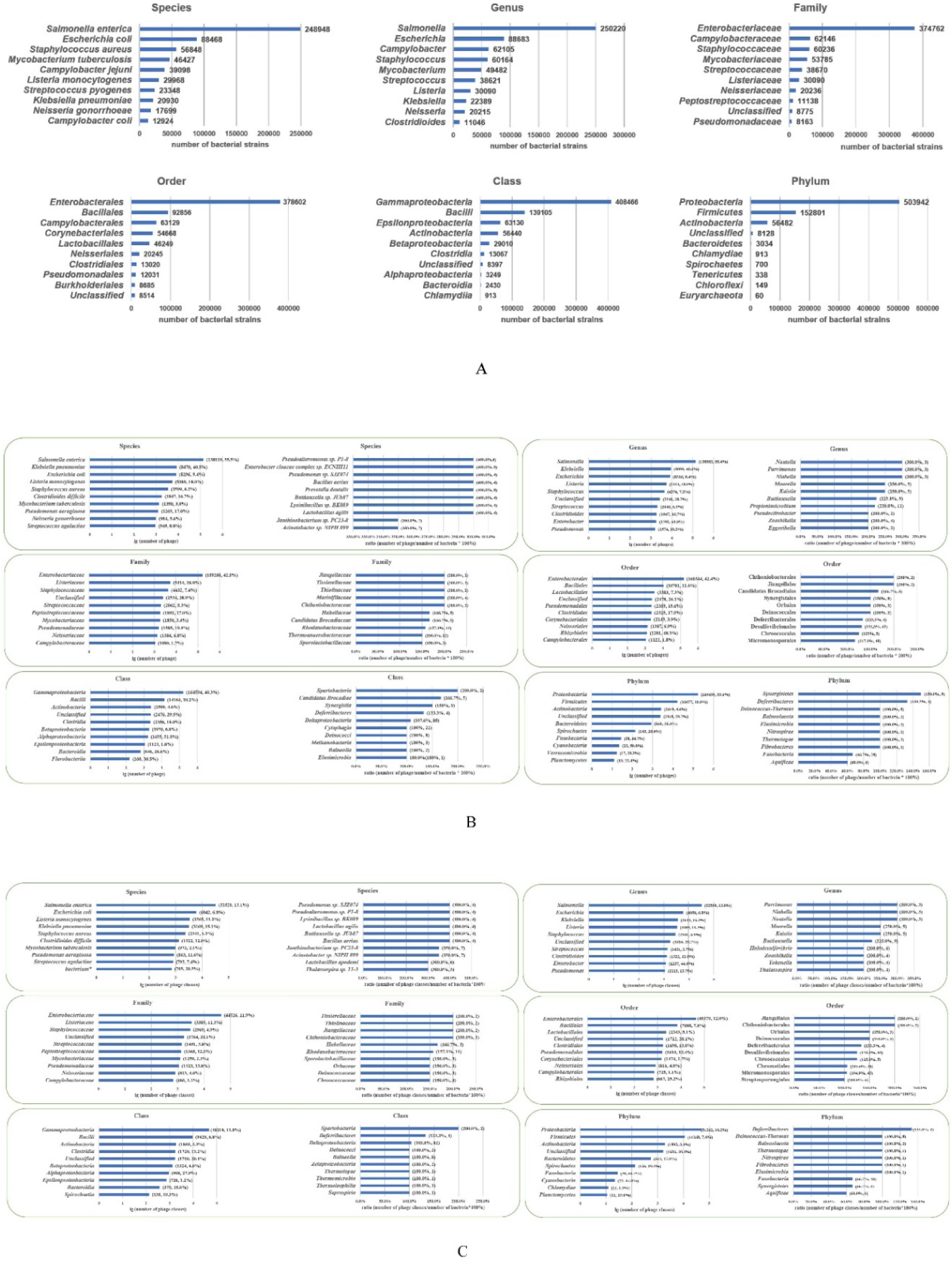
Number of complete temperate phage genomes and their hosts. All sp. species in the same genus are summed up in one species labeled as species sp., whose details can be found in Table. S3. (A) Top ten most bacteria at each taxonomic rank level. (B) Hosts that contain the top ten most temperate phages overall or per bacterium at each taxonomic rank level. (C) Hosts that contain the top ten most temperate phage entries overall or per bacterium at each taxonomic rank level.

**Extended Data Figure 4.**
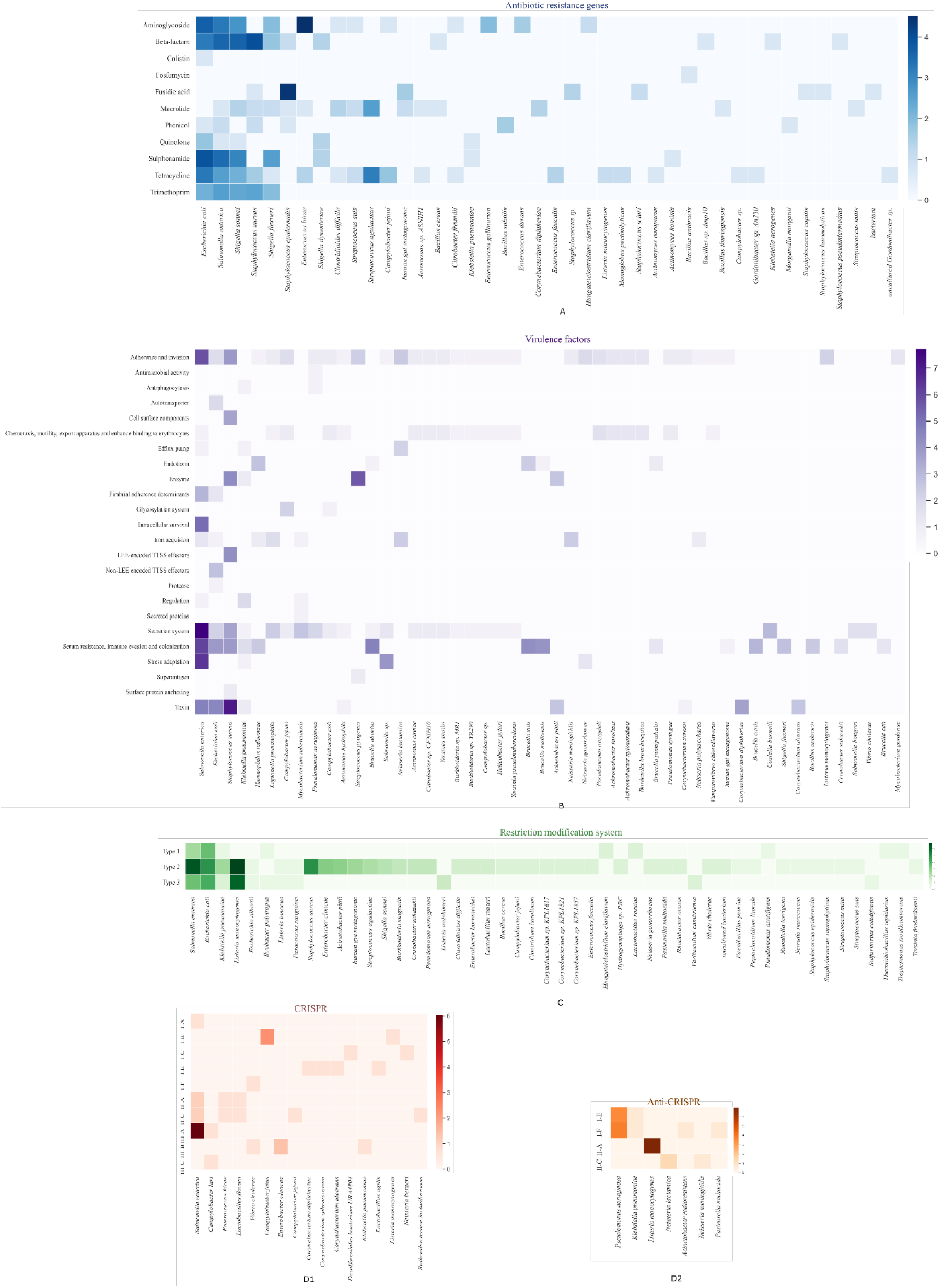
Heatmaps of the top 20 host species where their temperate phages contain the most antibiotic resistance genes, virulence factors, restriction-modification systems, CRISPR, and anti-CRISPR systems. (A) Heatmap of the top 20 host species where their temperate phages contain the most antibiotic resistance genes. Within one temperate phage, all the genes in the same gene type were counted as one. (B) Heatmap of the top 20 host species where their temperate phages contain the most virulence factors. Within one temperate phage, all the genes in the same virulence factor were counted as one. (C) The types of restriction-modification systems identified in temperate phages. The heatmap shows the top 20 temperate phages containing the most RM systems (classified by the host species of the phages). (D) The types of CRISPR and anti-CRISPR systems identified in temperate phages. (D1) the types of CRISPR systems identified in all temperate phages. (D2) the types of anti-CRISPR systems identified in all temperate phages.

**Extended Data Figure 5.**
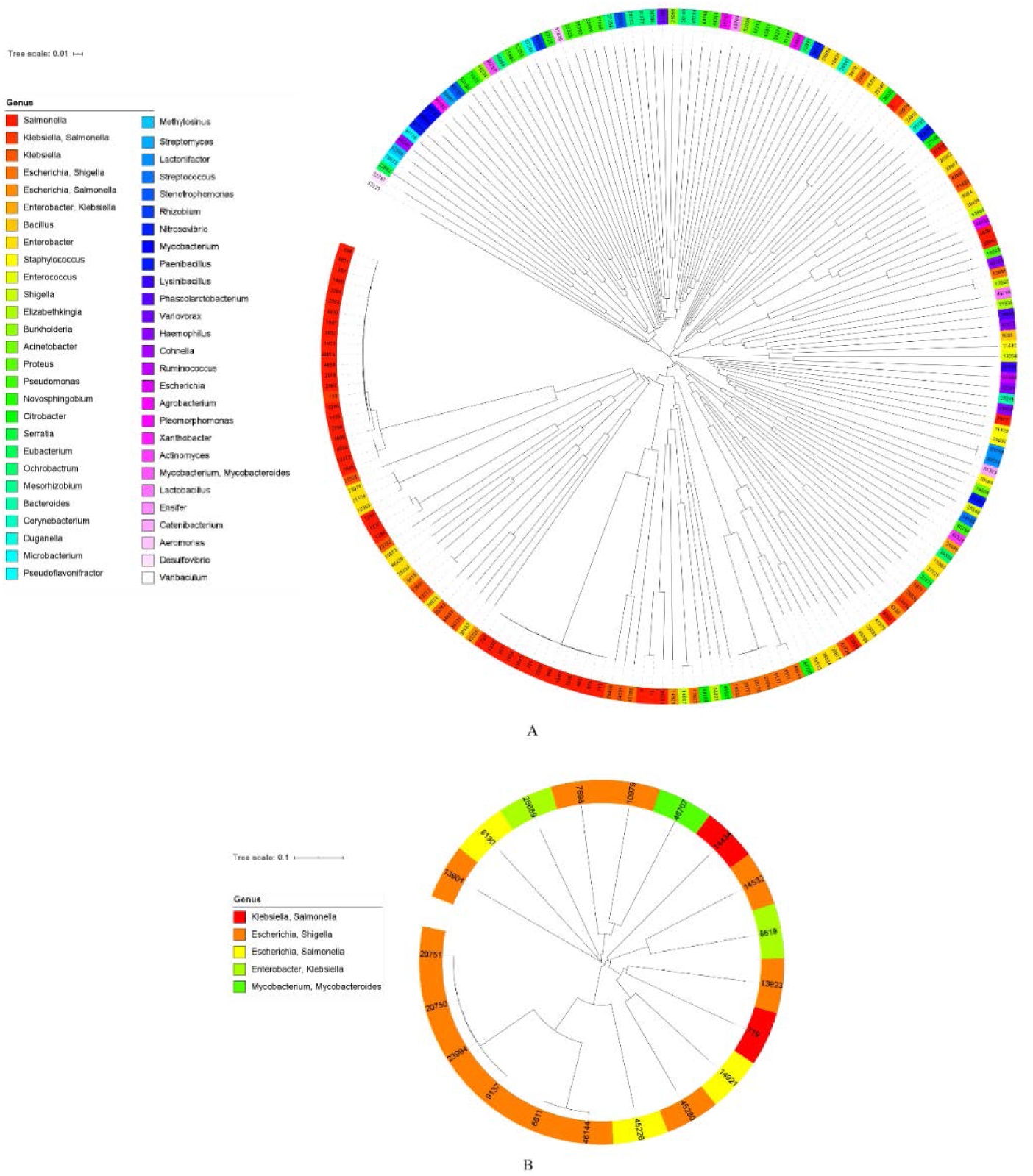
Host distribution and phylogenetic relationships of temperate phages infecting multiple hosts. (A) Phylogenetic relationships among phages with hosts at the level of genus. (B) Phylogenetic relationships among phages with hosts of cross-genera.

**Extended Data Figure 6.**
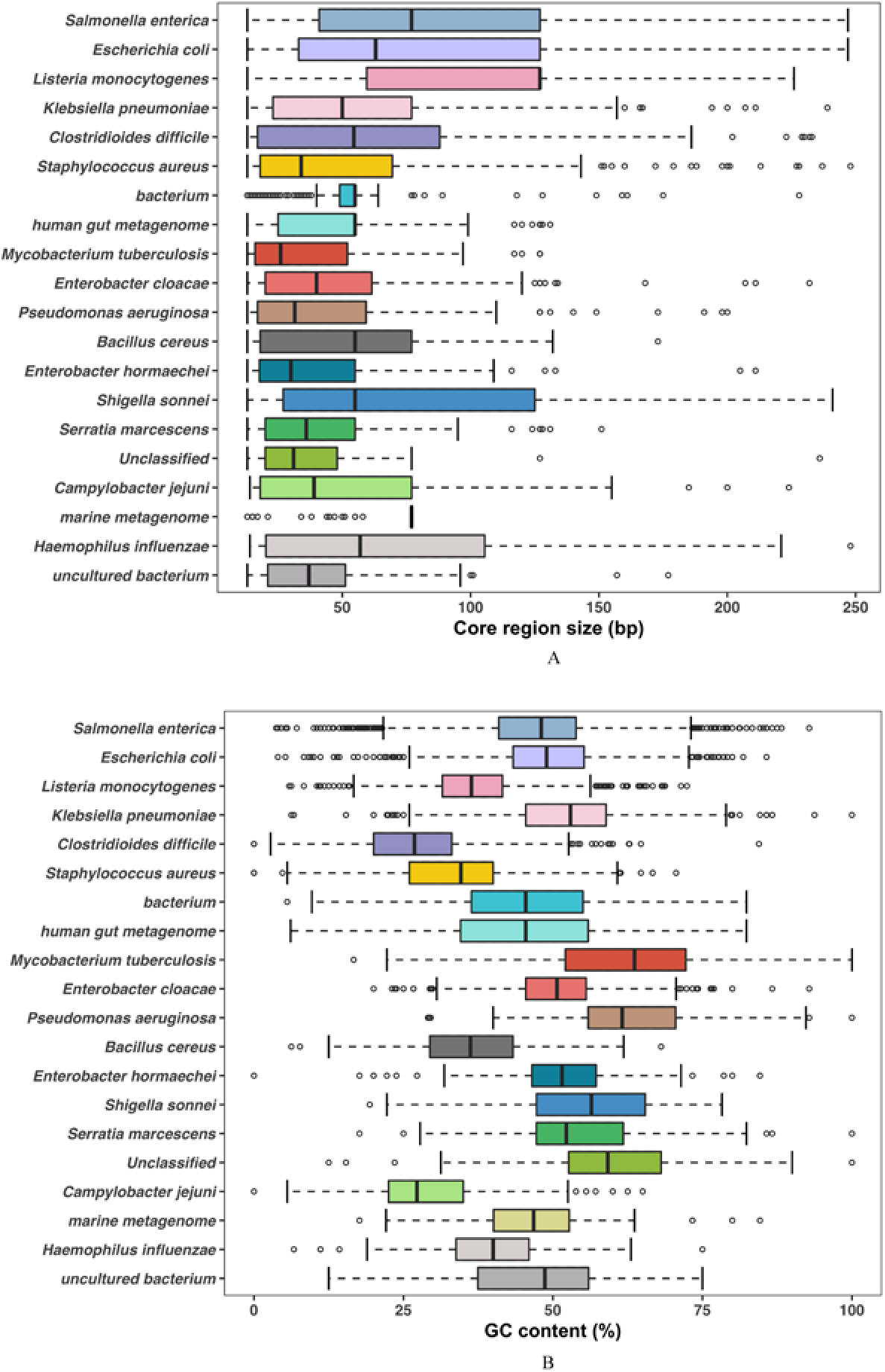
Genome size and GC content distribution of the core region of attachment sites in the top 20 host species. The distribution of core sequence size (A) and GC content (B) distribution in the top 20 most host species. The bacterium and Unclassified relates to the same name and NA listed in the species of GenBank taxonomy database, respectively. The ‘bacterium’ relates to the same name listed in the species of GenBank taxonomy database. ’Unclassified’ represents the original scientific name that cannot be classified into any species listed in the GenBank taxonomy database.

**Extended Data Figure 7.**
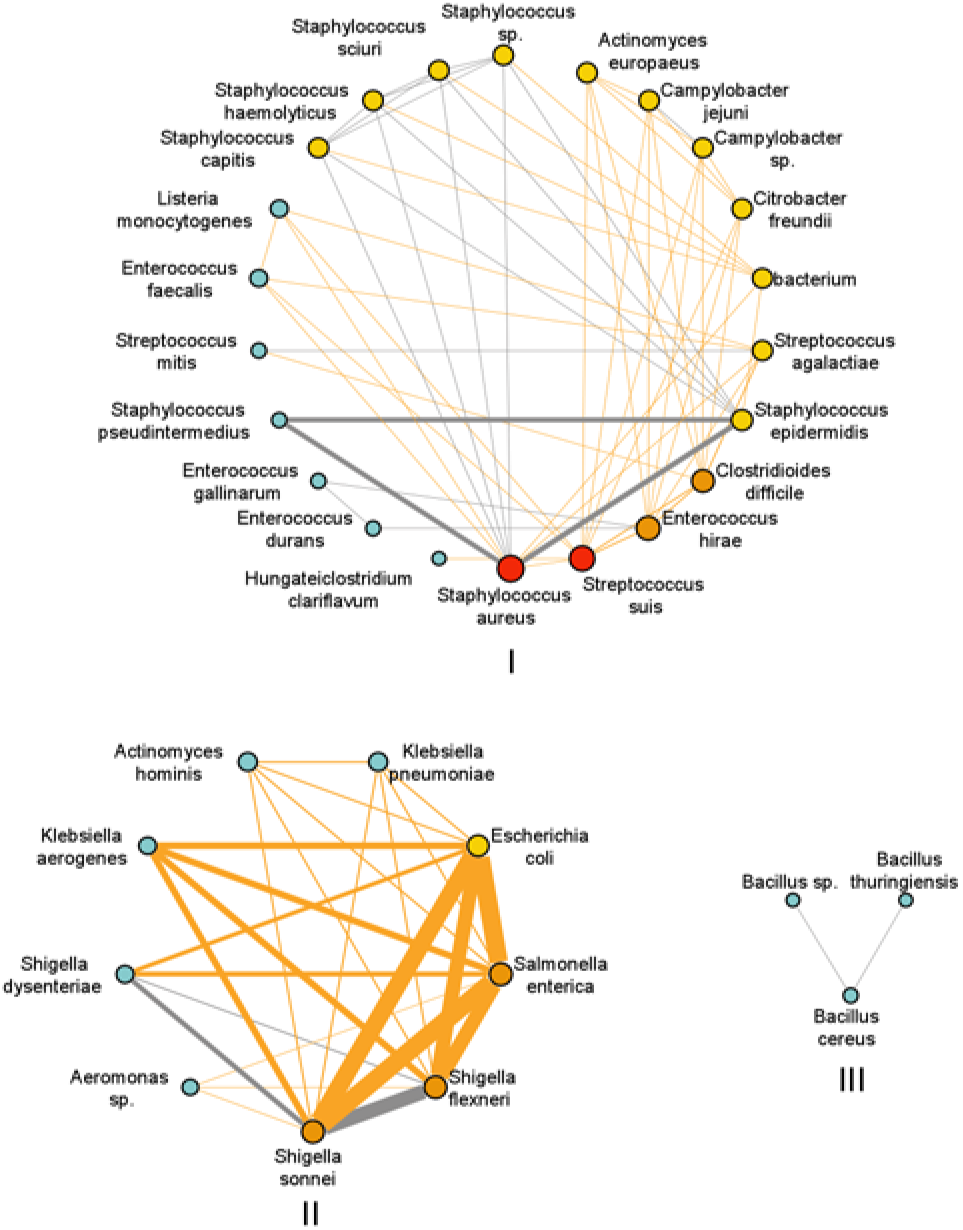
Connected network of antibiotic resistance gene (edge) within the temperate phages of different host species (nodes). The more the genes shared, the thicker the line and the darker the node is. The grey line represents the genes shared within the phages having the same host genus, while the yellow line represents the genes shared within the phages whose hosts were across genera.

**Extended Data Figure 8.**
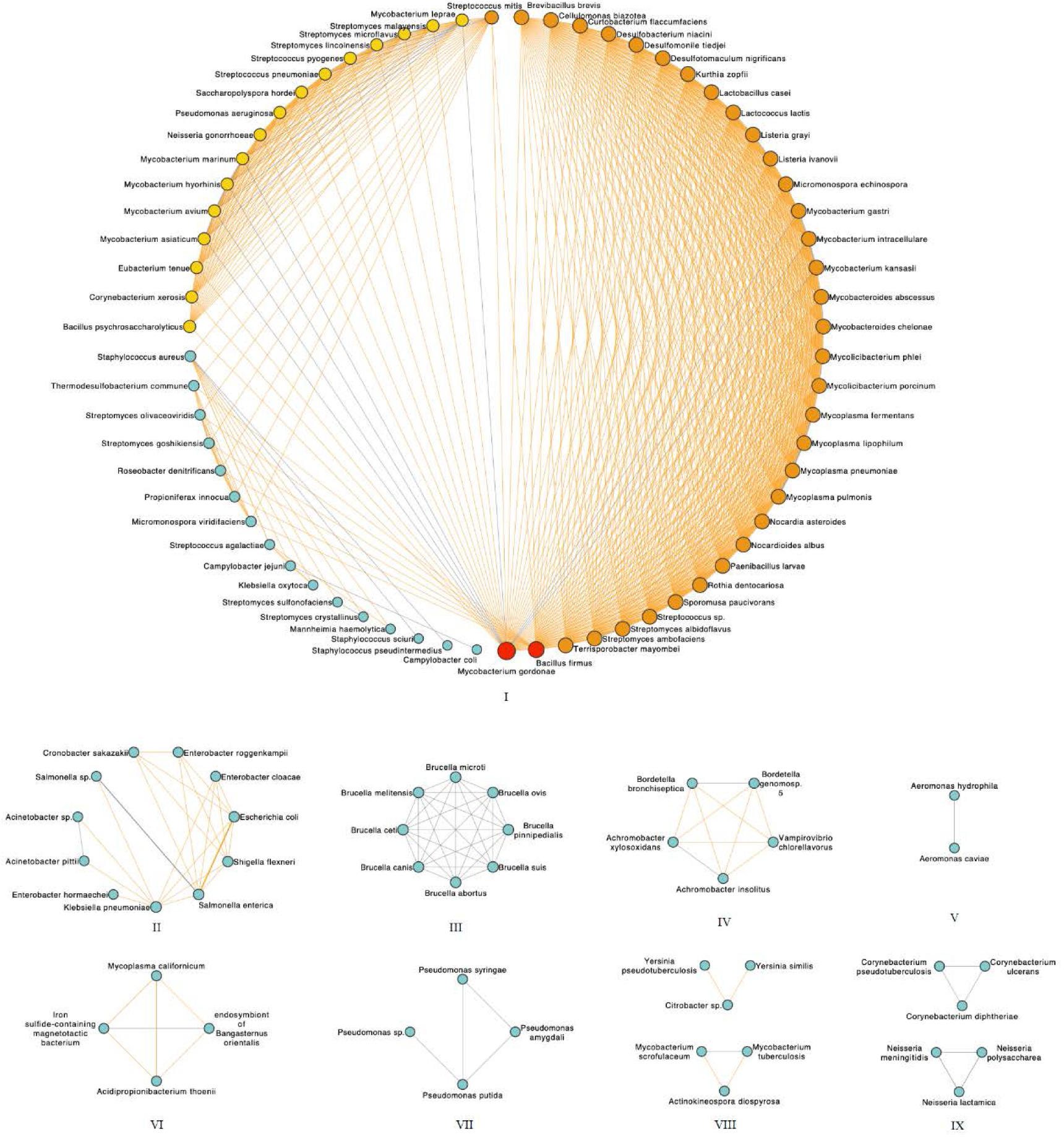
Connected network of virulence factor (edge) within the temperate phages of different host species (nodes). The more the genes shared, the thicker the line and the darker the node is. The grey line represents the genes shared within the phages having the same host genus, while the yellow line represents the genes shared within the phages whose hosts were across genera.

**Extended Data Figure 9.**
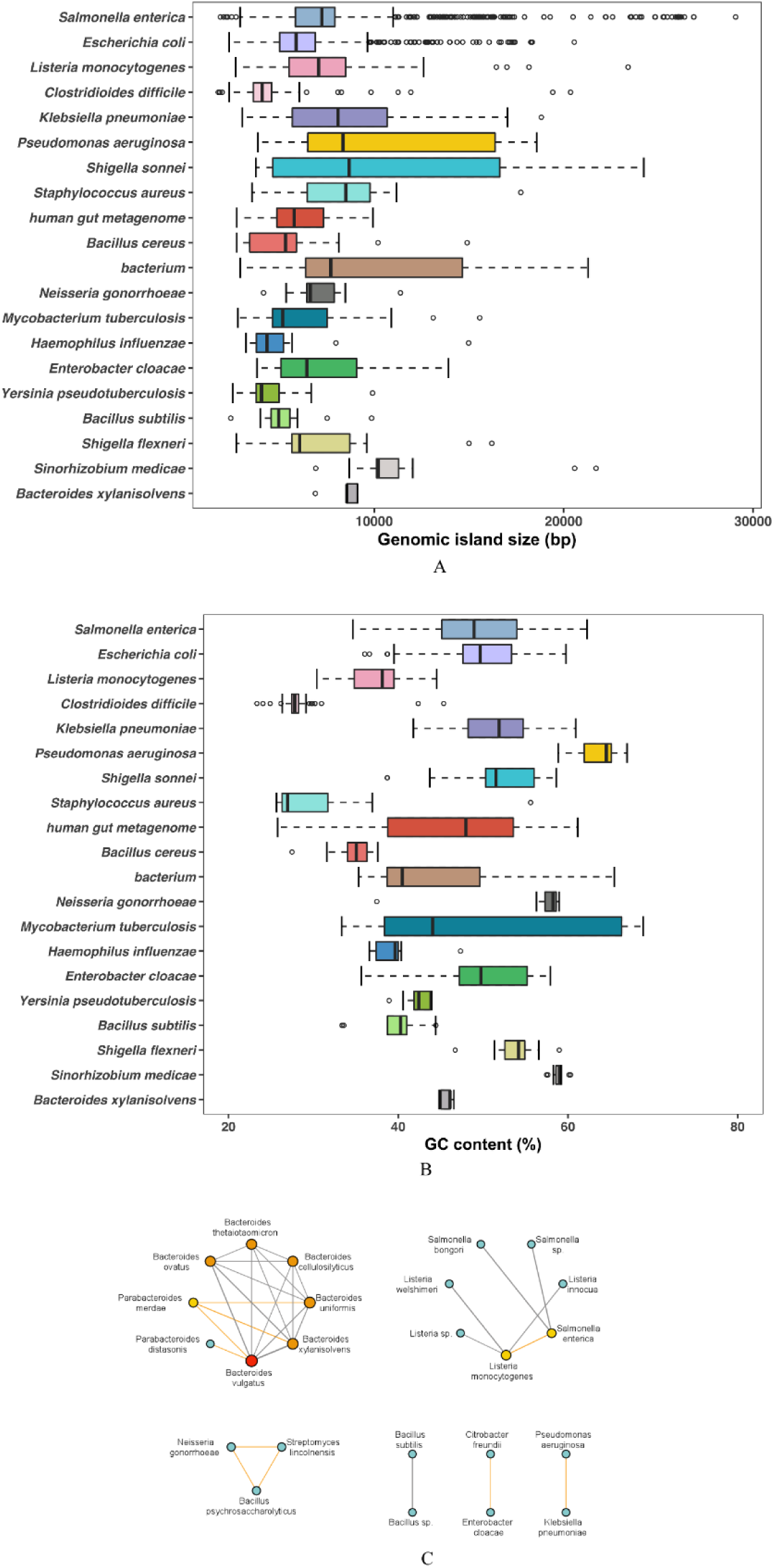
Genomic island (GI) distribution and sharing among temperate phages. The distribution of sequence size (A) and GC content (B) distribution in the top 20 most host species where their temperate phages containing the most GIs. (C) The connected network of GI (edge) within the temperate phages, where the GI was identified, in different host species (nodes). The more GI shared, the thicker the line and the darker the node is. The grey line represents the GIs shared within the phages having the same host genus, while the yellow line represents the GIs shared within the phages whose hosts were across genera.

**Extended Data Figure 10.**
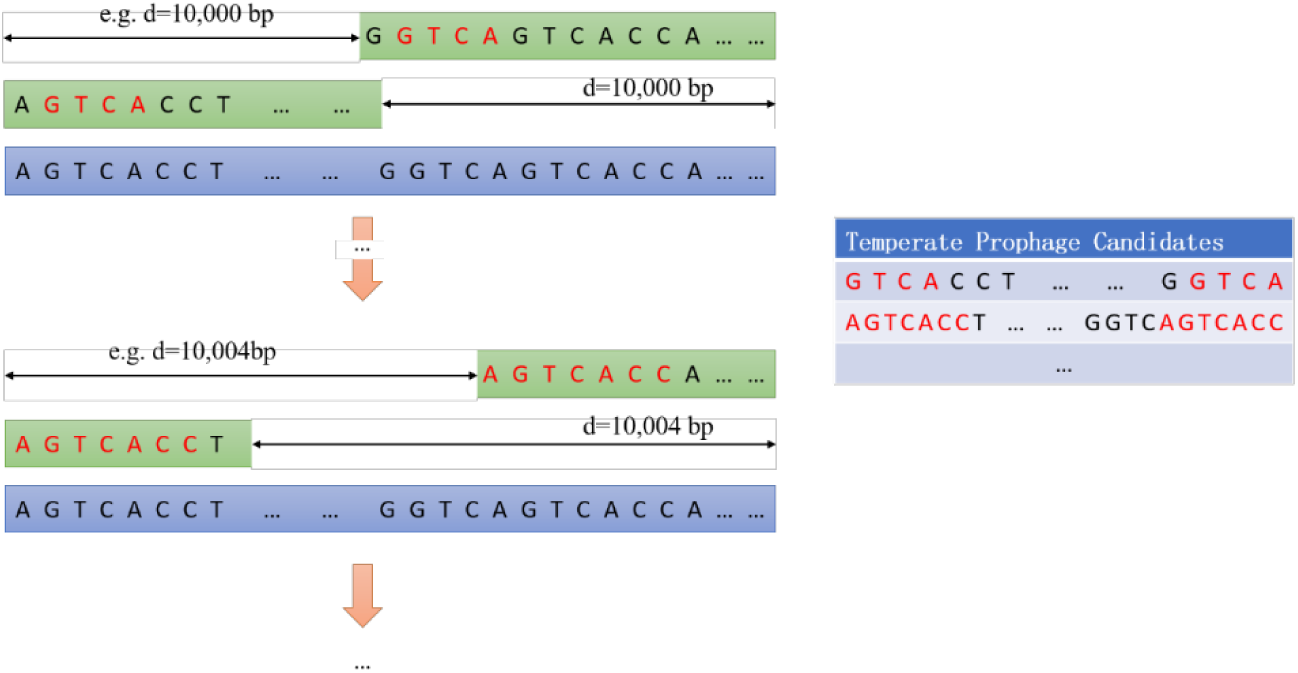
Search process of temperate phage candidates. The letters in red represent the repeats identified by our method.

**Extended Data Figure 11.**
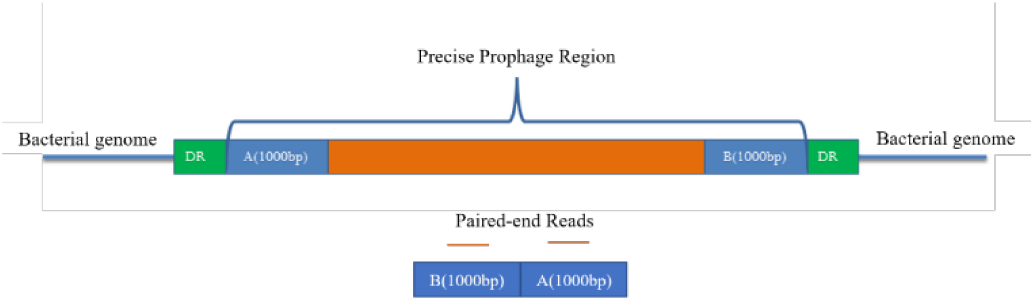
Verification of circular prophage sequence. DR represents directed repeat at both ends.

## Notes

https://github.com/NancyZxll/temperate-phage-active-prophage-detection.

https://phage.deepomics.org/

### Competing Interest Statement

The authors have declared no competing interest.

