## Supplemental Table 16 for "Mining bacterial NGS data vastly expands the complete genomes of temperate phages"

DBSCAN Algorithm

1: **Inputs:** W, E, M

2: **Output:** R

3: **DBSCAN(W,E,M)**

4: **Begin**

5: *Init* R←0

6: **for** each unvisited point y in W **do**

7: set y as visited

8: N←*getNeighbours(*y,E*)*

9: **if** *sizeOf(*N*)*<M **then**

10:set y as noisy data

11:**else**

12:R←next cluster

13:*ExpandCluster(*y,N,R,E,M*)*

14:**end if**

15:**end for**

16:**End**
